# Control of Craniofacial Development by the Collagen Receptor, Discoidin Domain Receptor 2

**DOI:** 10.1101/2022.02.03.479017

**Authors:** Fatma F. Mohamed, Chunxi Ge, Randy T. Cowling, Noriaki Ono, Abdul-Aziz Binrayes, Barry Greenberg, Vesa M. Kaartinen, Renny T. Franceschi

## Abstract

Development of the craniofacial skeleton requires interactions between progenitor cells and the collagen-rich extracellular matrix (ECM). The mediators of these interactions are not well-defined. Mutations in discoidin domain receptor 2 (DDR2), a non-integrin collagen receptor, are associated with craniofacial abnormalities, such as midface hypoplasia and open fontanels. However, the exact role of DDR2 in craniofacial morphogenesis is not known. As will be shown, *Ddr2*-deficient mice exhibit defects in craniofacial bones including impaired calvarial growth and frontal suture formation, cranial base hypoplasia due to aberrant chondrogenesis and delayed ossification at growth plate synchondroses. As established by localization and lineage tracing studies, *Ddr2* is expressed in progenitor cell-enriched craniofacial regions including sutures and synchondrosis resting zone cartilage, overlapping with Gli1+ cells, and contributing to chondrogenic and osteogenic lineages during skull growth. Tissue-specific knockouts further established the requirement for *Ddr2* in Gli1+ skeletal progenitors and chondrocytes and suggest important functions in chondrocyte proliferation and orientation as well as ECM organization. These studies establish a cellular basis for regulation of craniofacial morphogenesis by this understudied collagen receptor.

## Introduction

Craniofacial abnormalities affecting bone formation in the skull and face are common birth defects in humans.^1,2^ The craniofacial skeleton forms through regulated bone growth in the calvarium synchronized with growth of the developing brain and cranial base. Cranial sutures, fibrous tissues adjoining calvarial bones, are the principal sites for growth of the cranial vault. Sutures contain stem cells associated with several genetic markers including glioma-associated oncogene 1 (Gli1), an intermediate in hedgehog signaling, Axin-related protein 2 (Axin2), a component of the Wnt pathway, and paired related homeobox 1 (Prx1).^3–6^ The current paradigm suggests that sutural stem cells allow continued calvarial growth through osteogenic differentiation and bone formation at sutural edges of the calvarial bones. This process is coordinated with growth of the cranial base mediated by endochondral ossification centers called synchondroses. Histologically, the synchondrosis is a mirror-image growth plate with resting chondrocytes in the central zone and proliferative and hypertrophic chondrocytes on both sides. During endochondral ossification, the resting chondrocytes in the growth plate synchondrosis undergo sequential proliferation, hypertrophic differentiation, and apoptosis before being replaced by bone. The cranial base has been implicated in many human syndromes such as craniosynostosis (premature closure of cranial sutures), Down syndrome, Turner syndrome, cleidocranial dysplasia, cleft palate, and osteogenesis imperfecta.^7–11^

During craniofacial development, skeletal cells interact with the collagen-rich ECM through cell-surface receptors. Cell-matrix interactions regulate cell proliferation, migration, differentiation and remodeling to control and maintain shapes of individual cranial bones and the relative proportions of skeletal elements.^12^ Disruption of these interactions can cause craniofacial abnormalities.^12–15^ Discoidin domain receptors (DDR1 and DDR2) are a unique class of receptor tyrosine kinases (RTKs) that, together with β1 integrins, mediate interactions between cells and the ECM by specifically binding triple-helical collagens.^16–18^ DDRs contain a unique discoidin (DS) domain on their extracellular surface that is homologous to discoidin I-like domain of the slime mold *Dictyostelium discoideum*. DS domains are required for collagen recognition and specificity of DDRs such that DDR2 preferentially binds to fibrillar collagens I-III, but not nonfibrillar basement membrane type IV collagen that is exclusively recognized by DDR1. ^16,17,19^

*DDR2* plays an important role during human craniofacial morphogenesis, possibly being involved in skull changes occurring during the evolution of modern humans from archaic hominins^20^. Also, *DDR2* loss of function mutations cause the autosomal recessive growth disorder, spondylo-meta-epiphyseal dysplasia, with short limbs and abnormal calcifications (SMED, SL-AC). This disorder is associated with a number of craniofaciial abnormalities including a prominent forehead, open fontanelles, hypertelorism, a short nose with a depressed nasal bridge, long philtrum and micrognathia.^21–27^ *Ddr2*-deficient mice recapitulate craniofacial phenotypes seen in SMED, SL-AC patients including short snout and protruding eyes and display abnormal tooth development and impaired calvarial ossification.^28–31^ However, the basis for these abnormalities is not understood.

We recently described cell autonomous functions of *Ddr2* in growth of the appendicular skeleton^32^. Preferential localization of DDR2 in GLI1-positive skeletal progenitor populations associated with resting/growth zone chondrocytes of the growth plate and bone marrow was demonstrated while conditional inactivation of Ddr2 in *Gli1*-expressing cells and chondrocytes phenocopied the dwarf bone phenotype of globally *Ddr2*-deficient mice. In the present study, we focus on DDR2 functions in craniofacial bones. As will be shown, DDR2 is associated with putative suture stem cells that can serve as precursors for osteoblasts/osteocytes in the cranial vault and with synchondrosis-associated resting/proliferating chondrocytes that form hypertrophic chondrocytes, osteoblasts, and osteocytes in the cranial base. Furthermore, as shown by global and conditional inactivation approaches, DDR2 controls suture formation, frontal bone thickness and anterior/posterior growth of the skull. However, these DDR2 functions involve activity in different cell populations.

## Results

### Impairment in anterior-posterior skull growth and frontal suture/bone formation in *Ddr2*-deficient mice

To begin understanding the function of *Ddr2* in craniofacial development, we first defined which parts of skull are altered in global *Ddr2* deficiency using *Ddr2*^slie/slie^ mice. These animals contain a spontaneous 150 kb deletion in the *Ddr2* locus to generate an effective null^29^. We performed linear measurements on micro-CT scans of 3-month-old skulls (n=10) using previously described landmarks^33^. Our analysis along the anterior-posterior (AP) axis revealed a significant reduction (12%) in skull length (SL) mainly due to shortening of the nasal bone (NB), cranial vault (CV), and anterior and posterior cranial base (ACB and PCB) in *Ddr2*-deficient mice compared with WT littermates **(Fig. 1a-c)**. Accordingly, we detected a relative increase in skull height and width **(Fig. 1d-e)**. We further selected orthogonal planes in MicroView to measure the thickness of calvarial bones (**Fig. 1f**). Interestingly, frontal bone thickness was reduced by 55% with *Ddr2* deficiency; however, no significant differences were observed in the thickness of parietal or occipital bones (**Fig. 1f-g)**. Further analysis using Alcian blue and Alizarin red whole mount and H&E staining of 2-week-old skulls showed defective formation of frontal sutures in *Ddr2*-deficient mice (defect in 3/3 mice examined). In contrast, coronal and lambdoid sutures had a normal morphology (**Fig. 1h-i)**. *Ddr2*-deficient calvaria also had a reduced calvarial bone marrow cavity that was most prominent in the anterior skull bones **(Fig. 1f,i)**. No major differences were observed in transverse cranial sutures, such as coronal and lambdoid. Consistent with results from 2-week-old mice, reduced mineralization in the posterior frontal suture region was also seen in most skulls from 3-month-old *Ddr2*^slie/slie^ mice, although some phenotypic variability was observed at this age with 6/10 skulls affected **(Fig. 1b)**. In summary, skull deformities in *Ddr2*-deficient mice are associated with defects in AP skull growth, impaired frontal suture formation, frontal bone thickness and reduced calvarial bone marrow volume.

**Fig.1:**
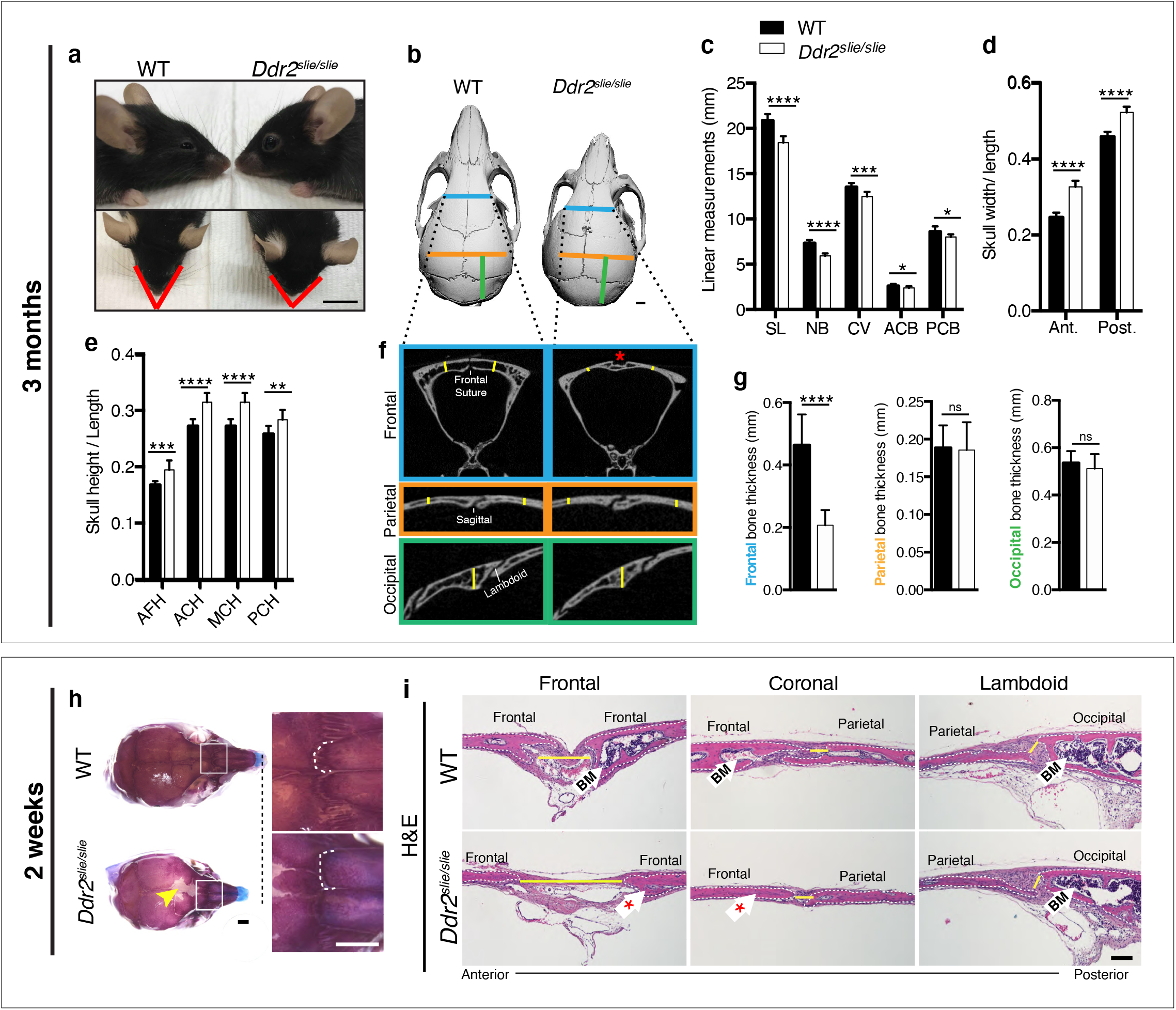
*Ddr2* deficiency results in impaired anterior-posterior skull growth with abnormal frontal bone and suture formation. WT and *Ddr2^slielsile^* mice were compared at 3 months **(a-g)** and 2 weeks **(h,i)**. **a-c**, Short snout, and reduced skull length in Ddr*2^slielsile^* mice. **a**, Side (upper) and top (lower) head views of 3 month-old *Ddr2^slie/slie^* mice and WT littermates. Scale bar: 1cm. **b**, 3D rendering of μCT scans of 3-month-old skulls. Scale bar: 1mm. **c**, Linear measurements along anteroposterior axis of skulls, where SL: skull length; NB: nasal bone; CV: calvaria vault; ACB: anterior cranial base; PCB: posterior cranial base. **d**, Quantification of anterior (ant.) and posterior (post.) skull width/length showed relatively increased skull width in *Ddr2^slie/slie^* vs WT mice. **e**, Linear measurements showed a relative increase in the anterior facial height (AFH) as well as anterior (ACH), middle (MCH), and posterior (PCH) cranial heights in *Ddr2^slie/slie^* skulls compared with WT. **f,g**, μCT scans of calvarial bones and quantification showing a significant reduction of frontal bone (in blue color) in 3-month-old *Ddr2^slie/slie^* mice in the absence of changes in parietal (orange) or occipital (green) calvarial bones. Note frontal suture defect in *Ddr2^slie/slie^* mice (red asterisk). Data are presented as mean ± SD. (*n*=10). \**P*<0.01 \*\**P*<0.01, \*\*\**P*<0.001, \*\*\*\**P*<0.0001, ns, not significant, two tailed- unpaired *t* test. **h**, Alcian blue and Alizarin red staining of 2-week-old mouse skull whole mounts shows delayed frontal suture formation and abnormal suture morphology (white dotted lines) in *Ddr2^slie/slie^* mice compared with WT. Boxed region is shown in higher magnification; right. Scale bar: 100μm. **i**, Hematoxylin and eosin (H&E) staining shows open frontal sutures in *Ddr2^slie/slie^*, but transverse sutures, such as coronal and lambdoid were not affected (highlighted by yellow lines). *Ddr2^slie/slie^* calvariae also had a smaller bone marrow cavity (Bm, white arrows) compared with WT. Frontal suture; coronal section; transverse sutures; sagittal section. Scale bar: 100μm.

### Reduced anterior-posterior skull growth in *Ddr2* deficiency is explained by cranial base hypoplasia due to abnormal chondrogenesis and delayed ossification of synchondroses

Skull elongation is mainly driven by endochondral ossification at the cranial base. The impaired AP skull growth observed in *Ddr2^slie/slie^* mice prompt us to examine the cranial base synchondroses and associated bones. We focused on intersphenoid (ISS) and spheno-occipital synchondroses (SOS) that form the anterior and posterior borders of basisphenoid bone (BS), the most affected cranial base bone in *Ddr2*^slie/slie^ mice (**Fig. 2c**). The BS bone forms the mid-posterior cranial base providing a foundation to support the brain, along with presphenoid (PS) and basio-occipital (BO) bones **(Fig. 2a)**. Our analysis showed that the ISS and SOS were abnormally wide in *Ddr2*^slie/slie^ mice compared with WT littermates (**Fig. 2a-d**). The width of the ISS and SOS was increased by 187% and 23%; respectively, while a significant increase in the height was only observed in the SOS of knockout mice (12%) (**Fig. 2b**). Accordingly, growth of basisphenoid bone was significantly inhibited in the knockout mice (reduced by 14%) (**Fig. 2c**). The basio-occipital (BO) bone was also significantly affected but to lesser extent. No major difference was seen in the presphenoid (PS) bone (**Fig. 2c**). H&E staining of 2-week-old skulls revealed that the *Ddr2*-deficient synchondroses, particularly ISS, showed severe defects in chondrocyte organization associated with loss of the columnar arrangement of proliferative chondrocytes (**Fig. 2d)**.

**Fig.2:**
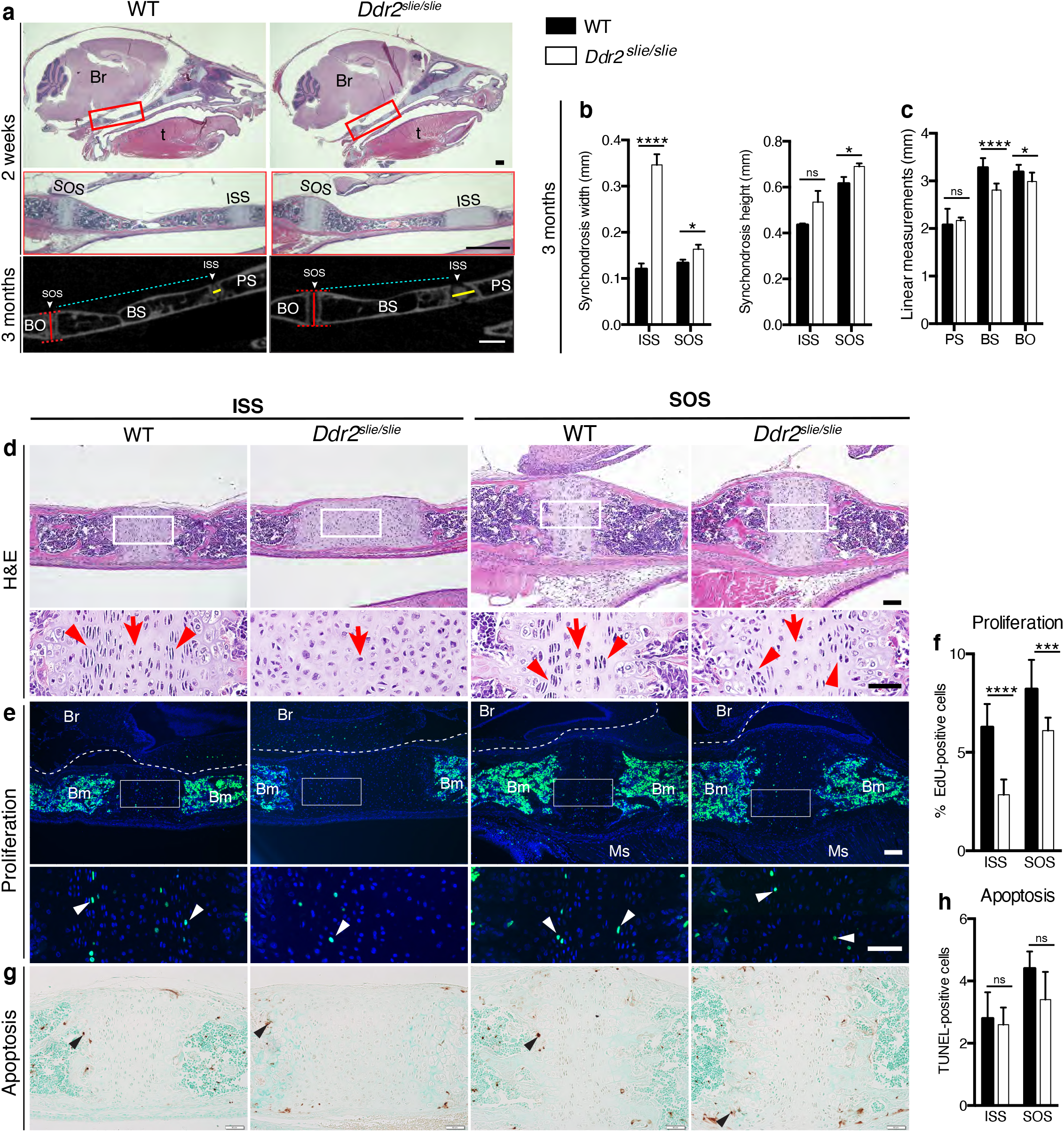
Cranial base hypoplasia due to chondrocyte disorganization and reduced chondrocyte proliferation in *Ddr2*-knockout synchondroses. **a**, H&E staining (upper) and μCT scans (lower) of WT and *Ddr2^slie/slie^* skulls showing wide cranial base synchondroses. Scale bar: 500μm. Boxed region (red) is shown in higher magnification. Br: Brain; t: Tongue. ISS: Intersphenoid synchondrosis; SOS: Spheno-occipital synchondrosis; PS: Presphenoid bone; BS: Basisphenoid bone; BO: Basis-occipital bone. In μCT scan of skulls (lower), arrowheads point to cranial base synchondroses; yellow and red lines highlight the width and height of synchondroses; respectively (Quantification is shown in **b**). cyan lines highlight shortening of basisphenoid bone between ISS and SOS in *Ddr2^slie/slie^* vs WT mice (Quantification of cranial base bone lengths is shown in **c**); Data are presented as mean ± SD. (*n*=10). \**P*<0.01, \*\*\*\**P*<0.0001, ns, not significant, two tailed- unpaired *t* test. **d**, H&E staining of ISS and SOS sections showing loss of columnar organization of proliferative chondrocytes (red arrowheads) in *Ddr2^slie/slie^* mice at 2 weeks of age. Red arrows point to resting chondrocyte zone. Boxed region is shown in higher magnification. Scale bar: 100μm. **e**, EdU staining (green) of ISS and SOS sections showing reduction in EdU+ cells (white arrowheads) in *Ddr2^slie/slie^* mice compared with WT littermates. Boxed region is shown in higher magnification. Scale bar: 100μm. Br: Brain, Ms: Muscle; Bm: Bone marrow. White dotted lines define the ventral surface of brain. **f**, Percentage of EdU+ cells in ISS and SOS of WT and *Ddr2^slie/slie^* mice. **g**, TUNEL staining (brown, black arrowheads) shows no changes in apoptotic levels between mice *Ddr2^slie/slie^* and WT. Cell nuclei were stained with methyl green (green). Scale bar: 50μm. **h**, Quantification of TUNEL-positive cells in cranial base synchondroses. Data in are presented as mean ± SD. \*\*\**P*<0.001, \*\*\*\**P*<0.0001, ns, not significant, two tailed- unpaired *t* test.

To begin to understand the cellular basis for the observed changes in the size of cranial base bones and synchondrosis structure, we assessed cell proliferation and apoptosis. EdU staining (shown in green) indicated labeling mainly in the proliferative chondrocyte zone of ISS and SOS (**Fig. 2e**, white arrowheads); however, EdU+ chondrocytes were significantly reduced in *Dd2*-deficient synchondroses: 55% in ISS and 26% in SOS, compared with WT controls (**Fig. 2f)**. We also performed TUNEL assay to measure apoptosis, but did not detect differences between *Ddr2*-deficient and control mice in either ISS or SOS regions (**Fig. 2g-h)**.

While a reduction in chondrocyte proliferation may explain the smaller BS and BO bones seen in Ddr2 deficient mice, proliferation changes do not readily explain the observed synchondrosis widening. To gain further insight into the pathogenesis underlying this phenomenon, we examined markers for resting and proliferative chondrocytes (type II collagen), hypertrophic chondrocytes (type X collagen) and for osteoblasts (type I collagen and integrin-binding sialoprotein (IBSP). In wild type mice, immunofluorescent staining of type II collagen was homogenously distributed in the cartilage matrix around chondrocytes. However, immunostaining was unevenly distributed in *Ddr2*-deficient synchondroses where more intense ring-like staining was observed around chondrocytes with diminished staining in the inter-territorial extracellular matrix (**Fig. 3a)**, indicating that DDR2 may regulate COL2 fibrillogenesis and/or ECM distribution. Type X collagen immunostaining showed a specific signal in the hypertrophic zone. However, we did not observe any change in the pattern of the type X collagen immunostaining between WT and *Ddr2*^slie/slie^ mice (**Fig. 3b)**. Immunofluorescence using COL1 and IBSP antibodies showed specific staining in the matrix of trabecular (Tb) and cortical (Ct) bones; however, staining intensity was reduced in *Ddr2*-deficient synchondroses compared with WT, suggesting defective bone formation and mineralization (**Fig. 3c,d)**. In summary, *Ddr2* deficiency is associated with abnormal chondrogenesis in the cranial base, disorganized chondrocytes, reduced chondrocyte proliferation, abnormal COL2 ECM distribution and delayed endochondral ossification.

**Fig.3:**
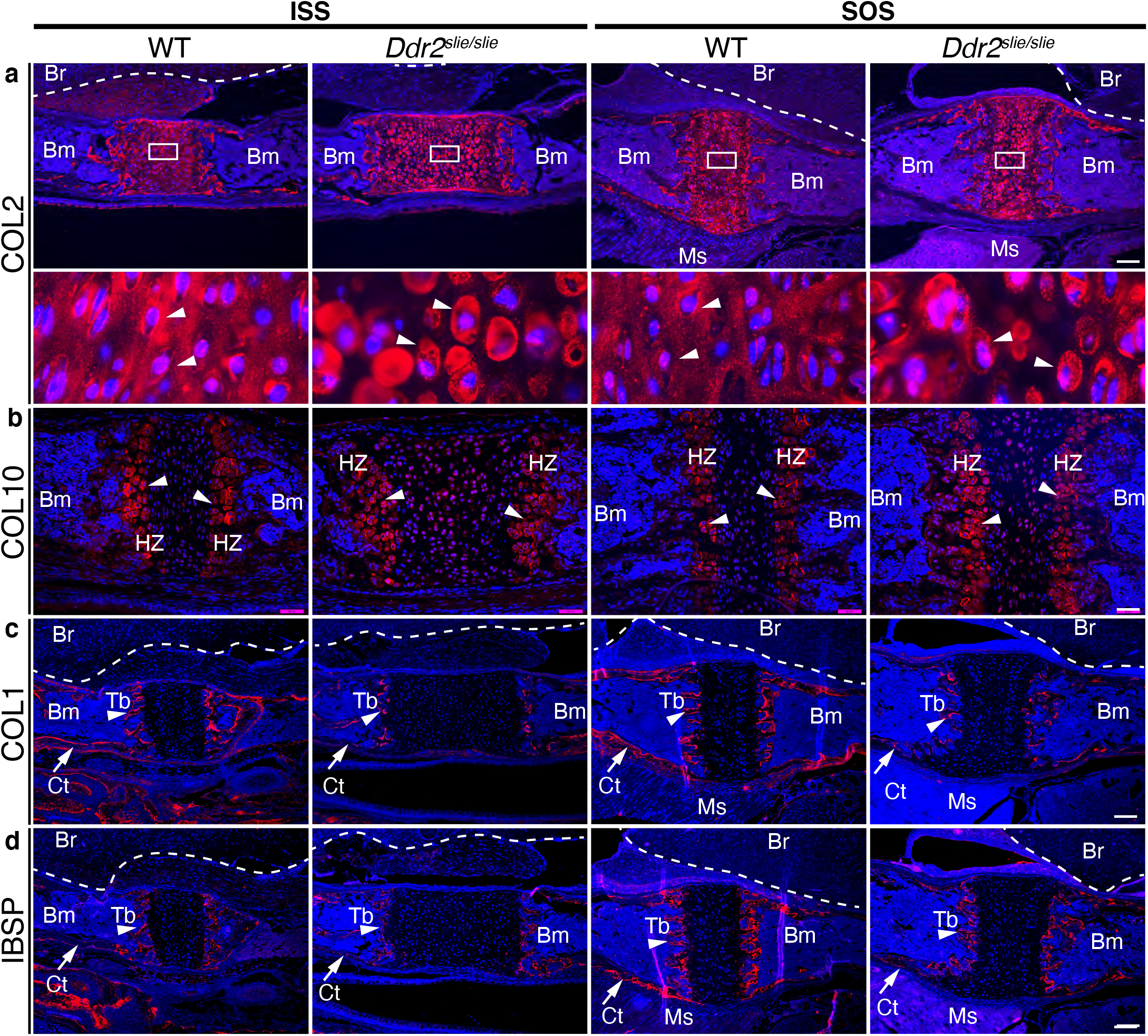
ECM defects in *Ddr2*-deficient synchondroses. **a-d**, Immunofluorescent staining of ISS and SOS sections from 2-week-old WT and *Ddr2^slie/slie^* synchondroses. **a**, Representative images of COL2 immunostaining (red) in ISS and SOS showing homogenous distribution around chondrocytes in WT synchondroses, while *Ddr2^slie/slie^* mice showed uneven, ring-like immunostaining around chondrocytes (white arrowheads). Boxed region is shown in higher magnification (bottom). Scale bar: 200μm. **b**, Immunofluorescent images of COL10 immunostaining in the hypertrophic zone (HZ) of synchondroses showing no major changes in staining distribution between WT and *Ddr2^slie/slie^* mice. Scale bar: 50μm. **c**, Immunofluorescence images showing COL1 (red) staining in trabecular (arrowheads) and cortical (arrows) bones of the cranial base in WT, but immunostaining is reduced in *Ddr2^slie/slie^* mice. Scale bar: 200μm. **d**, Immunofluorescent images showing IBSP (red) in trabecular (arrowheads) and cortical (arrows) bones of cranial base is decreased in *Ddr2^slie/slie^* synchondrosis compared with WT littermates. Scale bar: 200μm. Cell nuclei were stained with DAPI (blue) in **a-d**. Bm: Bone marrow; Br: Brain; Tb: Trabecular bone; Ct: Cortical bone; Ms: Muscle.

### Distribution of *Ddr2* expression in the craniofacial skeleton

To begin relating the cellular functions of DDR2 to the observed craniofacial phenotype of *Ddr2* deficient mice, we first examined the temporal and spatial distribution of *Ddr2*-expressing cells using a *Ddr2*-lacZ knock-in mouse model. The expression pattern of *Ddr2* was examined using a combination of whole mount and frozen sections from *Ddr2*^+/LacZ^ mice. During fetal development, LacZ staining was first detected at E11.5 (not seen at E9.5 -not shown). The whole mount staining at this time revealed broad *Ddr2* expression in the developing midface including the median and lateral nasal process (MNP and LNP), maxillary and mandibular processes, and around eyes (**Fig. 4a)**. A similar expression pattern was seen at E13.5 and E16.5 (**Fig. 4a)**. Using frozen sections of the E13.5 embryonic head, LacZ staining was detected in cartilage primordia of the cranial base, tongue mesenchyme, nasal septum, and Meckel’s cartilage in the developing mandible, and, as previously reported, developing tooth buds^31^. However, no LacZ staining was detected in the developing brain (**Supplemental Fig. 2a**).

**Fig.4:**
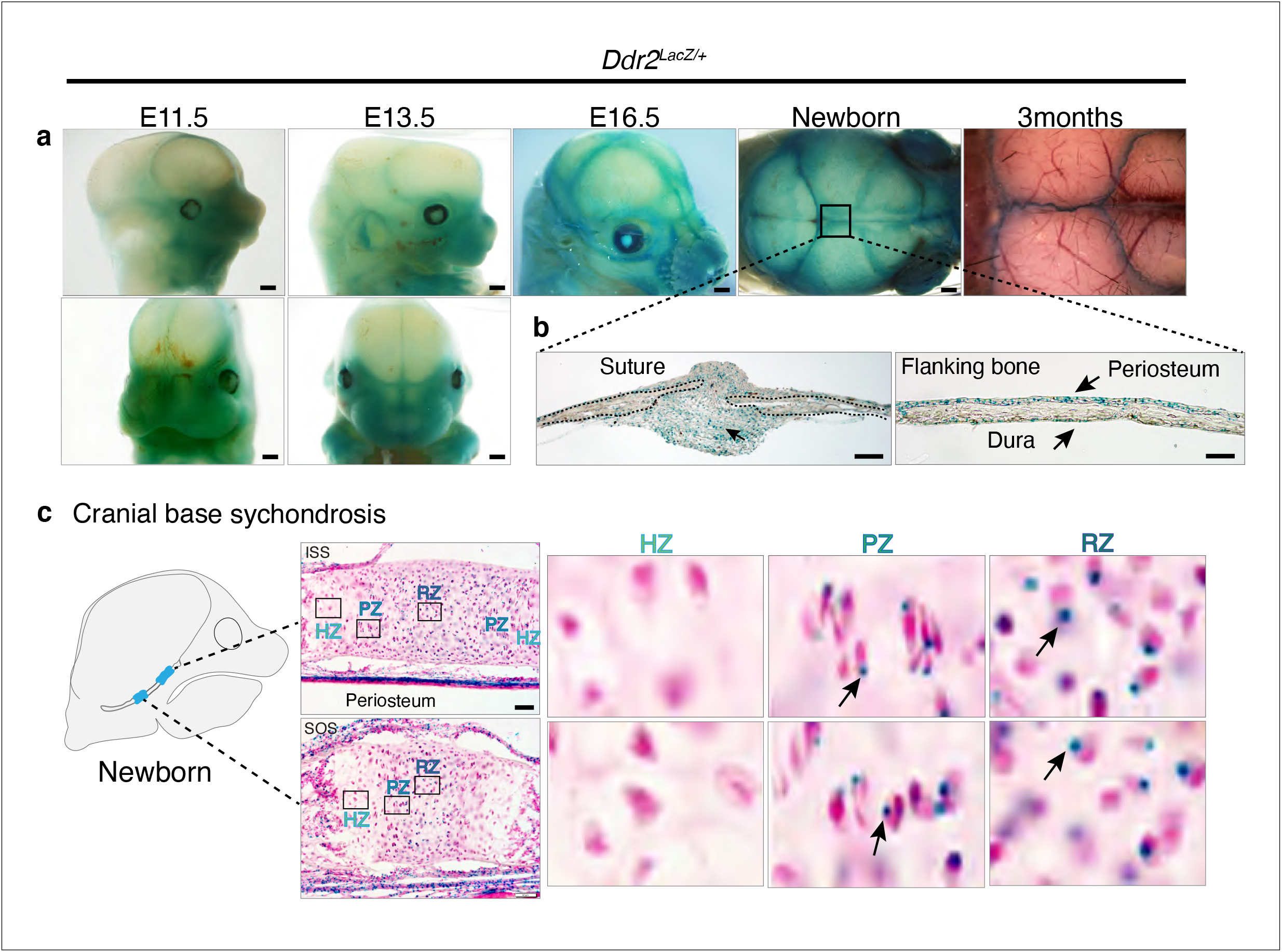
*Ddr2* expression in craniofacial skeleton. **a**, Whole-mount X-gal staining (green) of *Ddr2^Lacz/+^* skulls showing of *Ddr2* expression in midfacial region, cranial vault, and cranial sutures. Scale bar: 50μm. **b**, X-gal staining of cryostat sections of calvaria from newborn mice showing expression in suture mesenchyme, periosteum, and dura mater of flanking bones. Scale bar: 100μm, left and 50μm, right. **c**, X-gal staining of cryostat section of ISS (top) and SOS (bottom) from newborn mice revealing *Ddr2* expression in resting and proliferative chondrocyte zones, but low or undetected in terminal hypertrophic chondrocytes. Boxed regions are shown in higher magnification, right. Scale bar: 50μm.

Whole mounts of *Ddr2*^+/LacZ^ skulls from newborn mice showed broad *Ddr2* expression across the calvarial surface with preferential enrichment in cranial sutures. This distribution became more restricted to sutures in 3-month-old adult mice **(Fig. 4a)**. Frozen sections of neonatal *Ddr2*^+/LacZ^ skulls revealed *Ddr2* expression in the suture mesenchyme, periosteum of flanking bones, and dura mater on the ventral surface of calvaria **(Fig. 4b)**. Suture and periosteal expression were also seen in adult calvaria as well as in cells lining the bone marrow (**Supplemental Fig. 2b)**. However, LacZ staining was not seen in osteocytes, the terminally differentiated cells inside calvarial bones. This indicates that *Ddr2* expression is highest during early stages of osteoblast differentiation. *Ddr2* expression was also detected in synchondroses, primarily located in the resting (RZ) and proliferative chondrocyte (PZ) zone but low or undetected in terminal hypertrophic chondrocytes (HZ), and in the associated bone marrow and periosteum (**Fig. 4c)**. Overall, *Ddr2* expression was highest in regions enriched in skeletal progenitor cells including cranial sutures, periosteum and dura mater and resting chondrocytes. This suggest that *Ddr2* has functions in skeletal progenitor cells which contribute to development of the craniofacial skeleton.

### DDR2-expressing cells colocalize with GLI1 in calvaria sutures and cranial base synchondrosis where they contribute to osteogenic and chondrogenic lineages

Cranial sutures contain stem cells that contribute to craniofacial bone formation^6^. Given that *Ddr2* is expressed in sutures, we asked if *Ddr2* is in skeletal progenitor cells whose progeny can form the major cranial bone cell types. To address this question, we used a lineage-tracing approach by breeding heterozygous *Ddr2^Mer-icre-Mer^* female mice with male homozygous Ai14^CAG-tdTomato^ reporter mice. Cre-mediated recombination in *Ddr2^Mer-icre-Mer^*;Ai14^CAG-tdTomato^ mice was induced with intragastric tamoxifen injections given for 4 days after birth and then skulls were harvested and analyzed at day 5, 14, and 2 months of age (**Fig. 5a)**. Whole mounts of calvaria showed labeling that persisted for at least 2 months in all cranial sutures; frontal, sagittal, coronal and occipital sutures (**Supplemental Fig. 3a)**. At day 5, frozen sections revealed tdTomato labeling in a few cells in the cranial suture mesenchyme and in developing periosteum and dura mater of flanking calvarial bones (**Fig. 5b)**. At this age, the calvarial bone is very thin and devoid of a marrow cavity. Two weeks later, labeling was evident in cranial sutures, periosteum, and dura mater. Over a two-month chase period, labeling became intense in cranial suture mesenchyme, where undifferentiated cells reside, in the lining cells in the bone marrow of flanking calvarial bones, and in osteocytes (**Fig. 5b, Supplemental Fig. 3a)**. Therefore, the osteoblasts, osteocytes, and bone marrow-lining cells responsible for calvarial bone formation are derived from *Ddr2*-expressing cells.

**Fig.5:**
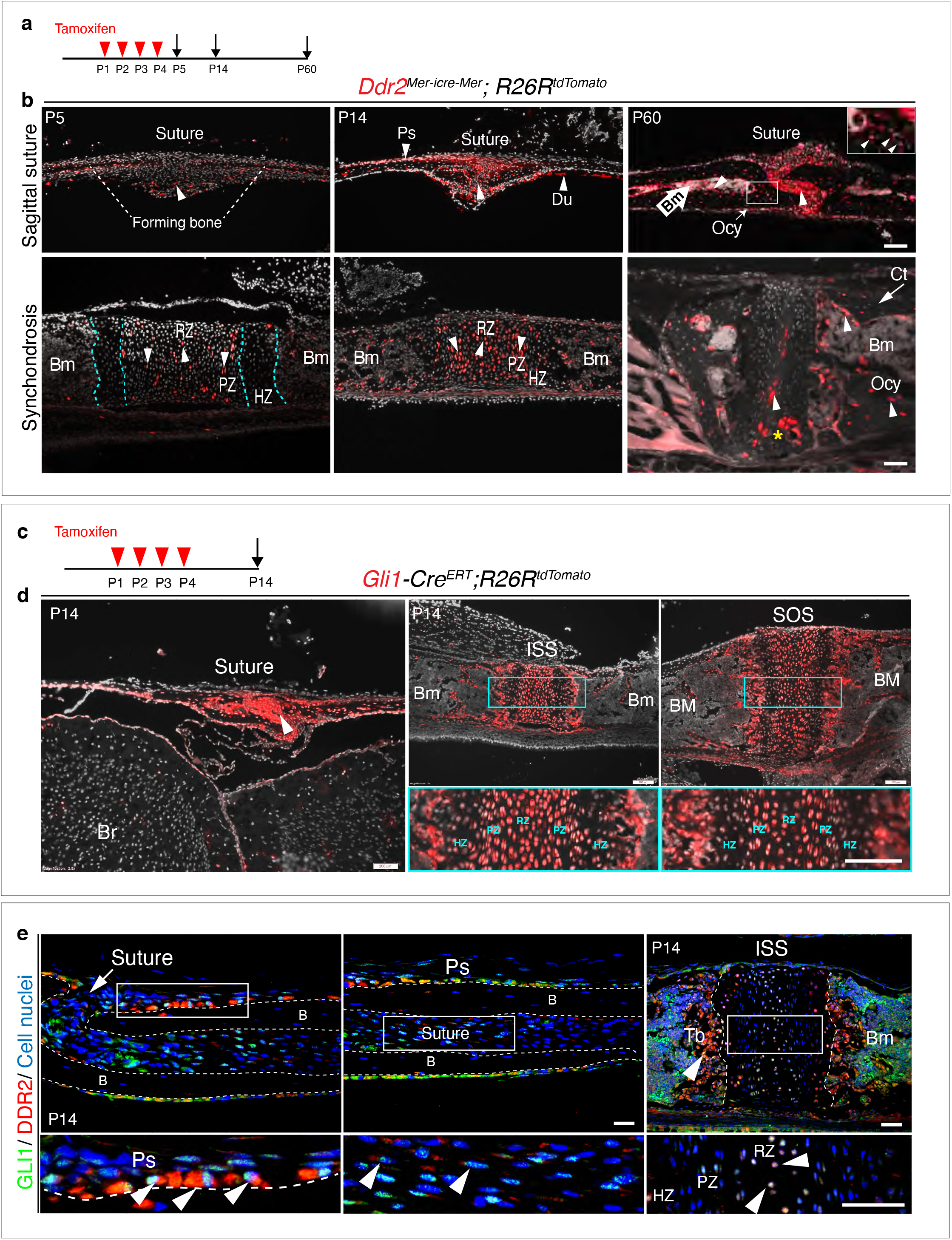
*Ddr2^mer-iCre-mer^* marks progenitors of the skeletal lineage during postnatal craniofacial development. **a**, Protocol used for induction of Cre-recombination and expression of tdTomato fluorescent protein (red) upon tamoxifen injection. **b**, Fluorescent tdTomato on cryostat sections of calvaria (upper panel) at the postnatal P5, P14 and P60 showing labeling (white arrowheads) in suture mesenchyme, periosteum (Ps) and dura mater (Dm), subsequently contributing to osteocytes (Ocy) (inset box) and bone marrow (Bm) of calvarial bones over time. Scale bar: 100μm (P14 and P60) and 50μm (P5). Lower panel, cranial base synchondrosis (ISS) showing labeling at P5 in resting (RZ) and proliferative (PZ) chondrocyte zones, but not in the hypertrophic zone (HZ) (cyan dotted lines), consistent with X-gal staining. Lineage trace at P14 shows an increase in tdTomato+ cells in all synchondrosis regions and associated bone marrow (Bm). At P60, tdTomato labeling is persistent in the middle zone and shows a clone of tdTomato labeling in proliferative and hypertrophic chondrocytes (yellow asterisk) and appears in the lining of bone marrow (Bm) and osteocytes (Ocy) of cortical bone (Ct). Scale bar: 100μm (P14 and P60) and 50μm (P5). Grey: cell nuclei. **c**, Protocol used for induction of Cre-recombination and expression of tdTomato fluorescent protein (red) in *Gli1-Cre^ERT^; R26R^tdTomato^* mice. **d**, Fluorescent tdTomato (red) on cryostat sections of calvaria (right) and cranial base synchondroses (ISS and SOS, left) from 2-week-old *Gli1-Cre^ERT^; R26R^tdTomato^* mice showing labeling in a similar cell population to that seen with *Ddr2^mer-iCre-mer^*. Scale bar: 200μm (Suture; left) and 100μm (ISS and SOS; right). Boxed region is shown in higher magnification (bottom). RZ: resting zone; PZ: proliferative zone; HZ: hypertrophic zone. **e**, Representative immunofluorescence images showing *Gli1* (green) and *Ddr2* (red) immunostaining of coronal sutures and ISS from 2-week-old mice. Scale bar: 100μm (ISS) and 50μm (suture). Boxed region is shown in higher magnification (bottom). White arrowheads indicate co-localization. Cell nuclei were stained with DAPI (blue). Ps: periosteum; B: Bone; Tb; trabecular bone

The cranial base synchondroses are a second major site of Ddr2 expression. TdTomato labelling was examined in both ISS **(Fig. 5b)** and SOS **(Supplemental Fig. 3b)**. At day 5, TdTomato labeling was detected in some cells in resting and proliferative zones, where *Ddr2*-labelled cells were either single cells dominant in the resting zone or pairs of daughter cells dominant in proliferative zone. However, *Ddr2* showed no labeling in hypertrophic chondrocytes. These results are in agreement with *Ddr2-LacZ* expression, indicating that tdTomato-labeled cells include *Ddr2*-expressing cells. By two weeks, progeny of *Ddr2*-labeled cells were seen in all chondrocyte lineages and associated bone marrow. By 2 months, labeled cells persisted in the resting chondrocyte zone and formed columns of cells extending through proliferative and hypertrophic zones. Progeny of *Ddr2*-expressing cells were also seen on the surface of trabecular bone, in the bone marrow and in osteocytes within the cortical bone of the cranial base (**Fig. 5b)**. Together, our results indicate that *Ddr2* is expressed in cell populations within cranial sutures and synchondroses having features of skeletal progenitor cells, but the identity of these cells still needs to be determined.

GLI1, a mediator of hedgehog signaling, is associated with stem/progenitor cells in cranial sutures.^6^ We asked if there is overlap between *Ddr2*- and *Gli1*-expressing cells. To address this question, we verified the activity of inducible Gli1-Cre^ERT^ under our experimental conditions by breeding with Ai14 td-tomato reporter mice. Newborn mice heterozygous for inducible Gli1 and the Ai14 allele were given four intragastric tamoxifen injections and analyzed after two weeks (**Fig. 5c)**. As previously reported,^6^ *Gli1-Cre^ER^* labeling was detected in cranial sutures in a pattern similar to that seen for *Ddr2* (**Fig. 5d)**. Gli1 also showed labeling in all chondrocyte lineages of cranial base synchondroses, with high concentration in resting and proliferative chondrocytes (**Fig. 5d)**. This suggests that Gli1 expression is not restricted to cells in cranial sutures or subchondral metaphyseal bone as previously reported, but it is also in growth plate chondrocytes of synchondroses. The similarity between *Ddr2* and Gli1 labeling in cranial sutures and synchondrosis suggests there may be a functional overlap between *Ddr2*- and *Gli1*-expressing cells. In support of this concept, we confirmed co-expression in the calvaria and synchondroses using co-immunostaining with DDR2 and GLI1 antibodies (**Fig. 5e)**. Colocalization was observed in select suture cells and adjacent periosteum and in the central resting zone of the ISS. The DDR2-GLI1 colocalization in synchondroses is reminiscent of what we previously observed in long bone growth plates where colocalization was preferentially seen in resting and proliferative zone chondrocytes ^32^.

### *Ddr2* functions in Gli1+ skeletal progenitors to control craniofacial morphogenesis

From the above results, we conclude that global *Ddr2* deficiency alters craniofacial morphology by affecting both calvarial ossification and frontal suture formation as well as endochondral bone growth in the cranial base. Furthermore, localization and lineage tracing studies suggest preferential expression of *Ddr2* in suture-associated skeletal progenitors and resting/proliferating zone chondrocytes in cranial base synchondroses. At least partial overlap was noted between DDR2 and cells expressing the skeletal progenitor marker, GLI1. *Ddr2* expression was also detected in resting and proliferating chondrocytes where type 2 collagen is present. To determine if Ddr2 has cell-autonomous functions in specific cell populations, a conditional deletion strategy was employed.

Our initial focus was on *Gli1*-expressing cells since this marker is associated with suture-associated skeletal progenitors as well as cranial base chondrocytes. *Gli1-Cre^ERT^* mice were crossed with mice carrying a floxed *Ddr2* allele, which includes exon 8 flanked by LoxP sites to generate conditional knockout mice (*Gli1-Cre^ERT^; Ddr2^fl/fl^* mice). To disrupt the *Ddr2* gene, we injected *Gli1-Cre^ERT^; Ddr2*^*fl*/fl^ newborn mice and their control littermates (*Ddr2*^*fl*/fl^) intragastrically with tamoxifen for four days and analyzed them at 3 months (**Fig. 6a)**. PCR analysis of genomic DNA extracted from ear punches confirmed the efficiency of recombination (**Fig. 6b)**. This was further validated by immunofluorescence of sections through the cranial base using an anti-pDDR2 antibody **(Fig. 6k)**. To investigate how *Ddr2* loss in *Gli1*-expressing cells affected the craniofacial skeleton, we performed AP linear skull measurements from 3-month-old mice as described in Figure 1. Tamoxifen-treated *Gli1-Cre^ERT^; Ddr2^fl/fl^* mice had craniofacial abnormalities of similar type and magnitude to those seen in global *Ddr2* knockouts. Conditional knockout mice had significantly reduced skull length (11%) due to shortening in the nasal bones, cranial vault, and posterior cranial base (**Fig. 6c-e)** and skull width and height were proportionately increased (**Fig. 6f-g)**. *Gli1-Cre^ERT^; Ddr2^fl/fl^* mice also had thin frontal bones (**Fig. 6d&h)**. Also, like *Ddr2*^slie/slie^ mice, delayed mineralization in the posterior portion of the frontal suture was observed in 5 of 10 mutant skulls; this was not seen in *Ddr2^fl/fl^* mice (n=10, **Fig. 6D**). Analyzing 2-week-old skulls, *Gli1-Cre^ERT^; Ddr2^fl/fl^* mice also had abnormally wide cranial base synchondroses with greatest changes seen in the ISS **(Fig. 6 i-k)**. *Gli1-Cre^ERT^* in the absence of the *Ddr2^fl/fl^* allele did not affect synchondrosis morphology **(Fig. 6i)**. Similarly, mutant mice showed a significant shortening in cranial base bones, including presphenoid (13%), basisphenoid (~15%), and basio-occipital bones (11%), suggesting impaired endochondral ossification at 3 months **(Fig. 6 m-p)**.

**Fig.6:**
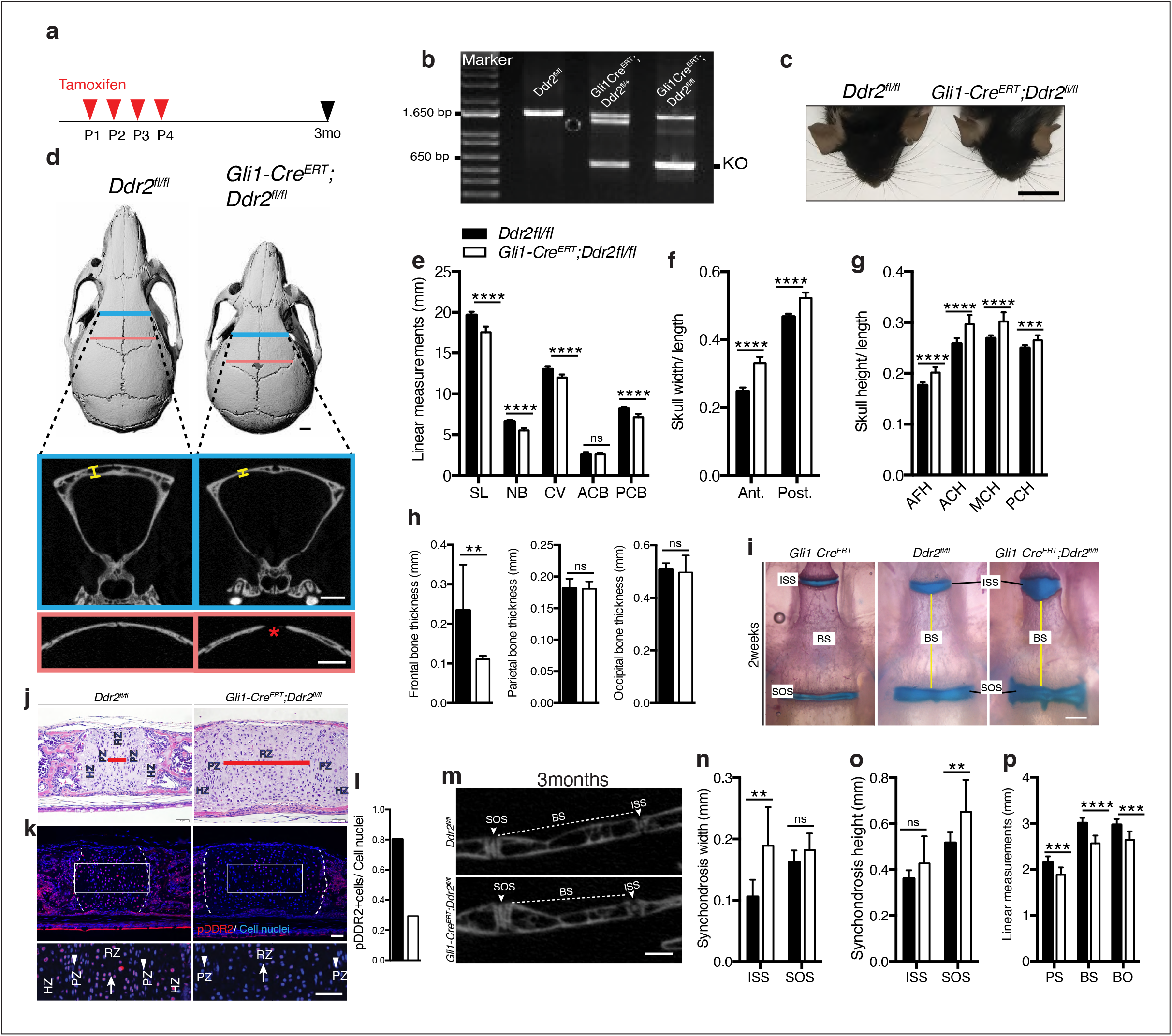
Loss of *Ddr2* in *Gli1*-expressing cells resulted in a craniofacial phenotype similar to *Ddr2^slie/slie^* mice. **a**, Protocol used for induction of Cre-recombination upon tamoxifen injection. **b**, Genotyping PCR showing WT (lower band) and Ddr2 floxed alleles (upper band) near 1,650 bp and recombined knockout allele below the 650 bp marker (KO). **c**, Top head view showing *Gli1-Cre^ERT^; Ddr2^fl/fl^* mice have short snout compared with *Ddr2^fl/fl^* mice. Scale bar: 1cm. **d**, μCT scans of *Ddr2^fl/fl^* and *Gli1-Cre^ERT^; Ddr2^fl/fl^* skulls show reduced anterior-posterior skull length and increased relative skull width and height (quantification in **e-g**). Note thinning and suture defect in the frontal bone in *Gli1-Cre^ERT^; Ddr2^fl/fl^* skulls. Scale bar: 1mm. **h**, Quantification of frontal, parietal, and occipital bone thickness. **i**, Alcian blue and Alizarin red whole mount staining shows *Gli1-Cre; Ddr2^fl/fl^* skulls have wide cranial base synchondroses compared with *Gli1-Cre and Ddr2^fl/fl^*. Scale bar: 500μm. **j**, H&E staining of ISS shows widening of resting zone and chondrocyte disorganization in *Gli1-Cre^ERT^; Ddr2^fl/f^* mice. Scale bar: 50μm. RZ: Resting zone; PZ: Proliferative zone; HZ: Hypertrophic zone. Red bar compares RZ width. **k**, Immunofluorescence images show reduced pDDR2 (red) immunostaining indicative of reduced DDR2 signaling in *Gli1-Cre^ERT^; Ddr2^fl/f^* synchondrosis. Dotted lines denote chondro-osseous junction. Boxed region is shown in higher magnification, lower panel. Cell nuclei were stained with DAPI (blue). Arrows, resting zone; arrowheads, proliferative zone. Scale bar: 50μm. **l**, quantification of immunostaining in **k**. **m-p**, μCT images and quantification show enlarged synchondroses associated with shortening in cranial base bone lengths in 3-month-old *Gli1-Cre^ERT^; Ddr2^fl/fl^* skulls compared with controls. Scale bar: 500μm. **c-h, m-p**, 3-month-old mice. **i-l**, 2-week-old mice. Data are presented as mean ± SD. (*n*=10). \**P*<0.01 \*\**P*<0.01, \*\*\**P*<0.001, \*\*\*\**P*<0.0001, ns, not significant, two tailed- unpaired *t* test.

Together, these results are consistent with the GLI1 distribution we observed in cranial sutures and synchondroses and suggest that DDR2 functions in GLI1-positive cells to both control suture formation/calvarial bone mineralization as well as endochondral growth in the cranial base. However, it is also possible that changes in suture formation/cranial vault growth could be secondary to reduced growth of the cranial base as has been previously proposed.^34^ If this were true, the cranial vault defects seen in *Ddr2^slie/slie^* and *Gli1-Cre^ERT^; Ddr2^fl/fl^* mice could be secondary to deficient growth at the cranial base rather than reflecting separate functions of *Ddr2* in sutures and calvarial bone. To discriminate between these possibilities, *Col2a1-Cre; Ddr2^f/f^* mice were developed to selectively deleted *Ddr2* in chondrocytes.^35^ At birth, mutant mice were viable and indistinguishable from their control littermates, but as they matured, *Col2a1-Cre; Ddr2^fl/fl^* mice exhibited growth defects compared with littermate controls (*Ddr2^fl/fl^*). The *Col2a1-Cre* transgene by itself did not cause any change in the cranial base synchondroses as demonstrated by whole mount staining (**Fig. 7i)**. Inhibition of *Ddr2* signaling in cranial base cartilage was confirmed by pDDR2 immunofluorescence (**Fig. 7j)** as well as PCR analysis of DNA extracted from ear cartilage (not shown). Linear measurements on micro-CT scans of 3-month-old skulls (n=10) showed significantly reduced AP skull growth and increased skull width and height in *Col2a1-Cre; Ddr2^fl/fl^* mice (**Fig. 7a-d)**. However, the reduction in AP skull length (7%) did not reach levels seen in *Ddr2*^slie/slie^ (12%) or *Gli1-Cre^ERT^; Ddr2^fl/fl^* skulls (11%) (**Fig. 1 versus 6)**. *Col2a1-Cre; Ddr2^fl/fl^* mice also exhibited growth defects in endochondral bones, mainly BS and BO contributing to cranial base hypoplasia **(Fig. 7h)**. In addition, the ISS was significantly wider (270%) and higher (32%) (**Fig. 7g,f)**. The width and height of the SOS in *Gli1-Cre^ERT^; Ddr2^fl/fl^* was also increased, but to a lesser extent than than for the ISS **(Fig. 7g)**. This phenotype is like that seen in *Ddr2*^slie/slie^ and *Gli1-Cre^ERT^; Ddr2^fl/fl^* mice. In contrast, conditional deletion of *Ddr2* in chondrocytes minimally affected the AP growth of cranial vault (decreased by 2.5%) (**Fig. 7b)**. *Col2a1-Cre; Ddr2^fl/fl^* mice exhibited significant thinning in the frontal bone (56%) as compared with controls (**Fig. 7a,e)**. Unlike *Ddr2^slie/slie^* or *Gli1-Cre^ERT^; Ddr2^fl/fl^* mice, the parietal bone (11.4%,) and occipital bone (12%) also showed a moderate decrease in thickness-11.4 and 12%, respectively **(Fig. 7e)**. Significantly, conditional knockout in Col2a1-Cre lineages did not affect cranial sutures which were indistinguishable from *Ddr2^f/f^* controls (n=10 mice/genotype, **Fig. 7a)**. This clearly resolves functions of *Ddr2* in cranial base that were disrupted in *Ddr2^slie/slie^, Gli1-Cre^ERT^; Ddr2*^*fl*/fl^ and *Col2a1-Cre; Ddr2^fl/fl^* mice from functions in sutures that were only disrupted in *Ddr2*^slie/slie^ and *Gli1-Cre^ERT^; Ddr2^fl/fl^* animals and establishes separate functions for *Ddr2* in both suture mesenchyme and cranial base chondrocytes.

**Fig.7:**
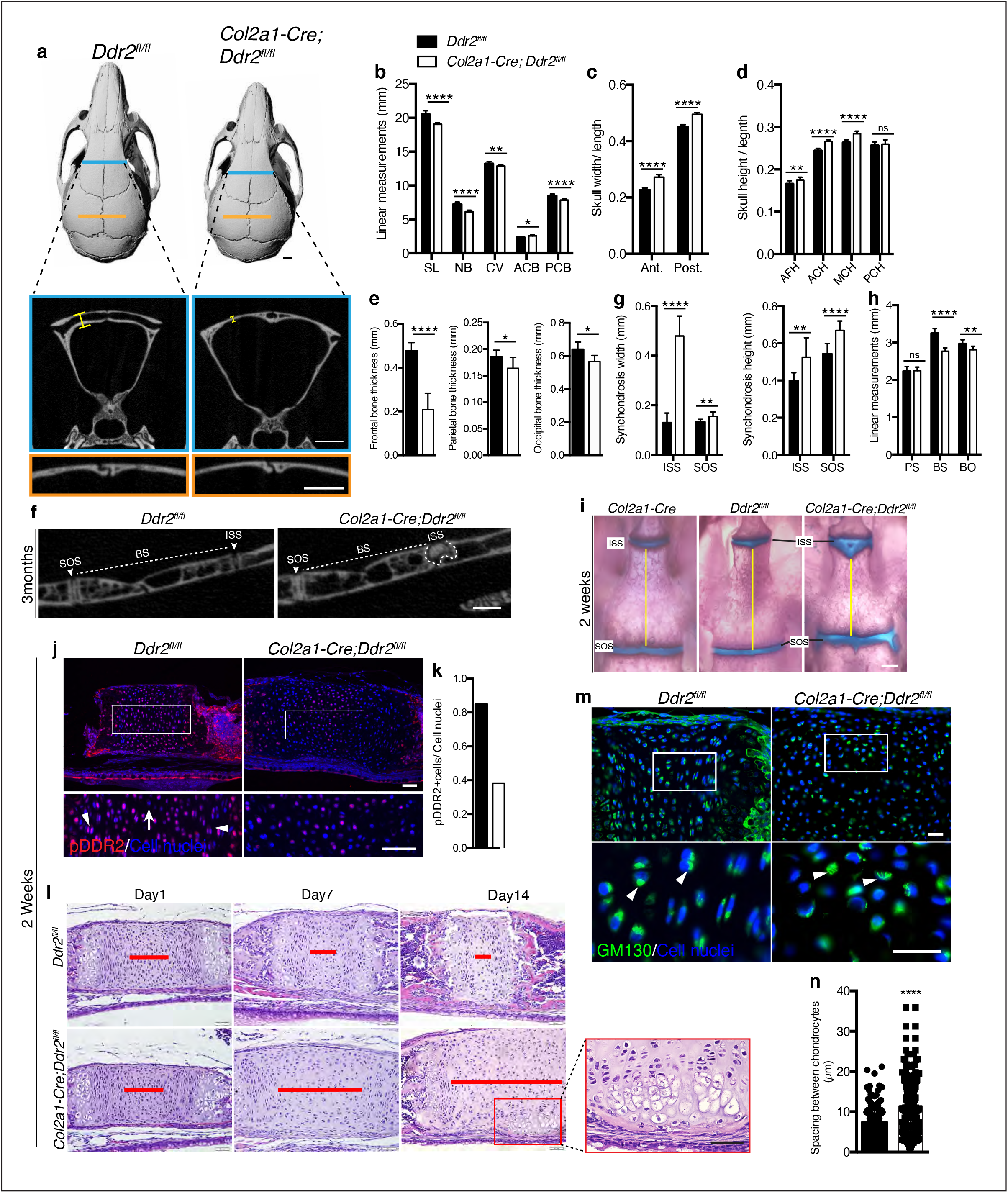
*Ddr2* conditional knockout in *Col2a1*-expressing chondrocytes causes cranial base hypoplasia and alters synchondroses without affecting cranial sutures. **a**, μCT scans of *Ddr2^fl/fl^* and *Col2a1-Cre; Ddr2^fl/fl^* skulls showing reduced anterior-posterior skull length, length of individual bones and increased relative skull width to height in conditional knockout mice (quantification in **b-d**). Note thinning of frontal bone in *Col2a1-Cre; Ddr2^fl/fl^* skulls, but no effect on cranial sutures. Scale bars: 1mm. **e**, Quantification of frontal, parietal, and occipital bone thickness. **f-h**, Quantification using μCT scans showing enlarged synchondroses associated with shortening in cranial base bone lengths in *Col2a1-Cre; Ddr2^fl/fl^* skulls. Data are presented as mean ± SD. (*n*=10). \**P*<0.01 \*\**P*<0.01, \*\*\**P*<0.001, \*\*\*\**P*<0.0001, ns, not significant, two tailed- unpaired *t* test. **(i)** Alcian blue and alizarin red whole mount staining showing *Col2a1-Cre; Ddr2^fl/fl^* skulls had wide cranial base synchondroses compared with *Col2a1-Cre and Ddr2^fl/fl^*. Scale bar: 500μm. **j**, Immunofluorescence images showing reduced pDDR2 (red) immunostaining indicating loss of DDR2 signaling in *Col2a1-Cre; Ddr2^fl/fl^* mice (Quantification in **k**). Scale bar: 50μm. Boxed region is shown at higher magnification, lower panel. Cell nuclei were stained with DAPI (blue). **l-n**, Analysis showing *Col2a1-Cre; Ddr2^fl/fl^* mice exhibited time-dependent widening in resting zone, altered polarization and ectopic hypertrophy. **l**, Time-course analysis using H&E staining shows no difference in histological structures of the ISS between *Col2a1-Cre; Ddr2^fl/fl^* mice and their control littermates at P1, but during the first two weeks, the resting zone became abnormally wide (red lines) and exhibited ectopic hypertrophy on the ventral side of cranial base synchondrosis (red box). Scale bar: 50μm. **m**, Immunofluorescent images of GM130 staining (green) shows well-defined Golgi staining adjacent to the nucleus of cells in RZ of wild type synchondroses, but in mutant synchondroses, GM130 immunostaining is diffuse and ill-defined indicating disturbed cell organization. Boxed region is shown in higher magnification, lower panel. Cell nuclei were stained with DAPI (blue). Scale bar: 20μm. **n**, Linear measurements shows increased spacing between chondrocytes in resting zone of mutant synchondroses. Spacing between chondrocytes was measured by drawing lines between chondrocytes in resting zone using ImageJ. **a-h, m-p**, 3-month-old mice. **i-k, m,n**, 2-week-old mice. Data are presented as mean ± SD. (*n*=3). \*\*\*\**P*<0.0001, two tailed-unpaired *t* test.

In contrast to results with *Gli1-Cre^ERT^; Ddr2*^*fl*/fl^ and *Col2a1-Cre; Ddr2^fl/fl^* mice, conditional deletion of *Ddr2* in mature osteoblasts using *OC-Cre* did not affect skull length, cranial sutures, or the cranial base in 3-month-old mice (**Supplemental Fig. 4**). This Cre line induces efficient recombination in mature osteoblasts/osteocytes^36^ and we detected efficient excision of the floxed *Ddr2* allele in DNA from vertebrae-containing tail biopsies from *OC-Cre;Ddr2^fl/fl^* mice (result not shown). These results are consistent with craniofacial functions of *Ddr2* being restricted to skeletal progenitors and chondrocytes rather than mature bone forming cells.

### Loss of *Ddr2* in chondrocytes causes ectopic hypertrophy and disrupted cell polarization

The presence of abnormal synchondroses and associated cranial base hypoplasia was a consistent finding after global or conditional *Ddr2* inactivation. In all cases, reduced growth of BS and BO bones was observed and chondrocyte organization into resting, proliferative and hypertrophic zones was disrupted. To further define the onset and evolution of synchondrosis changes in *Ddr2* deficiency, we conducted a time-course histological analysis focusing on ISS morphology of *Col2a1-Cre; Ddr2^fl/fl^* mice **(Fig. 7I)**. At birth, no significant differences in synchondroses were seen between mutant and control littermates. However, as endochondral ossification progressed, the resting zone (highlighted with red lines) became progressively narrower in controls while expanding at the expense of the proliferative and hypertrophic zones in Col2a1-Cre; Ddr2^fl/fl^ mice (**Fig. 7l)**. As was the case in *Ddr2^slie/slie^* mice **(Fig. 2)**, synchondrosis widening could not be explained by increased cell proliferation which was reduced in ISS and SOS (**Supplemental Fig. 5a**). The question became why mutant synchondroses were enlarged and exhibited an increase in the cell number, particularly in the central resting zone (**Fig. 7l**). It is possible that chondrocytes proliferating at a lower rate were retained in the resting zone rather than transitioning to the proliferative zone. As was seen in *Ddr2^slie/slie^* mice, type II collagen distribution was abnormal in *Col2a1-Cre; Ddr2^fl/fl^* mice with a selective loss of staining in the inter-territorial matrix that was accompanied by an increase in average spacing between cells (**Supplemental Fig. 5c, Fig. 7n**). Together, these changes contributed to the observed widening in the synchondrosis resting zone.

Intriguingly, P14 *Col2a1-Cre; Ddr2^fl/fl^* mice formed an ectopic hypertrophic zone on the ventral side of the ISS at right angles to the normal synchondrosis axis (**Fig. 7l)**. MicroCT images of 3-month-old mice also showed W-shaped synchondroses in *Col2a1-Cre; Ddr2f^l/fl^* mice (**Fig. 7f**), suggesting ectopic endochondral ossification taking place in the ventral side of the ISS. This phenotype was seen in 8 out of 10 mutant mice examined. With the shift in chondrocyte organization, we asked if cell polarity was disrupted in the absence of *Ddr2*. To test this, we examined the cellular distribution of GM130, a Golgi apparatus marker that normally maintains a fixed orientation to the nucleus in chondrocytes^37^. In control mice, immunostaining of GM130 showed well-defined, localized staining adjacent to the nucleus, indicating that positional information was retained within the growth plate (**Fig. 7m)**. In mutant synchondroses; however, GM130 immunostaining was disorganized, suggesting that chondrocyte polarization was disrupted. This was also seen in *Gli1-Cre^ERT^; Ddr2^fl/fl^* synchondroses (data not shown). Together, our findings indicate that proper DDR2 signaling is important for specification of cell polarity possibly through interaction with the cartilage extracellular matrix; however, a direct role of *Ddr2* on chondrocyte polarity cannot be excluded.

## Discussion

The ECM is a critical regulator of skeletal development and has been implicated in a wide spectrum of human skeletal disorders, many of which involve craniofacial structures.^12–15^ In this study, we highlight the importance of collagen-cell interactions with emphasis on cell-autonomous functions of the cell surface collagen receptor, DDR2, during growth of the craniofacial skeleton. Characterization of craniofacial phenotypes of globally *Ddr2*-deficient mice revealed two types of defects; 1) abnormalities in development of the flat bones of the skull including defective frontal suture formation/mineralization and thinning of frontal bones, 2) reduced anterior-posterior skull growth related to delayed endochondral ossification at cranial base synchondroses. To understand the basis for these defects, we conducted a systematic characterization of the distribution and lineage of *Ddr*2-expressing cells and used a conditional deletion approach to resolve the cellular sites of action of *Ddr2*. These studies complement our previous analysis of DDR2 function in development of the appendicular skeleton^32^.

As a prerequisite for understanding function, we determined the spatial distribution of *Ddr2* expression during craniofacial development. Analysis of LacZ knock-in reporter mice revealed expression in developing midface as early as E11.5 with staining initially seen in cartilage primordia of the cranial base, nasal septum, and Meckel’s cartilage in developing mandible. Later during development, *Ddr2* expression was localized in neonatal calvariae in developing periosteum, dura mater and cranial sutures where craniofacial stem cells reside. In the adult, *Ddr2* was also localized to cranial sutures, periosteum, dura mater and in bone marrow lining cells. *Ddr2* expression was either low or undetected in osteocytes. *Ddr2* was also localized to the resting and proliferative zones of neonatal cranial base synchondroses but was not detected in hypertrophic chondrocytes. These findings led us to hypothesize that *Ddr2* regulates development of the craniofacial skeleton through its function in skeletal progenitor cells and chondrocytes.

This concept is further supported by lineage tracing studies conducted in *Ddr2^Mer-icre-Mer^;Ai14^CAG-tdTomato^* mice where an initial tamoxifen pulse labelled cells in cranial suture mesenchyme, periosteum and dura as well as resting and proliferating cells of synchondroses. Over a two-month chase period, the TdTomato label increased in sutures and expanded into lining cells of bone marrow and calvarial osteocytes while, in the cranial base, labelled cells persisted in the central resting zone and expanded to proliferative and hypertrophic zones of synchondroses and into adjacent bone. While these studies do not prove *Ddr2 i*s expressed in stem/progenitor populations, they show DDR2-positive cells can serve as precursors to cells of the bone lineage. More detailed studies involving longer-term lineage tracing as well as isolation and characterization of *Ddr2*-expressing cells will be required to determine if this gene is active in a self-renewing skeletal progenitor population.

Consistent with DDR2 functioning in skeletal progenitors, we observed substantial colocalization of DDR2 with GLI1 in calvarial sutures, periosteum and synchondroses. GLI1-positive sutural stem cells (SuSCs) are of great interest because of their contribution to the development of craniofacial bones and possible role in craniofacial abnormalities. Ablation of GLI1+ SuSCs leads to craniosynostosis, a premature closure of cranial sutures resulting in arrested skull growth,^6,38^ while impaired cell proliferation and osteogenic differentiation of SuSCs is associated with wide-open fontanels due to reduced or delayed bone growth.^39,40^ To determine if DDR2 functions in GLI1+ cells, we conditionally inactivated *Ddr2* using *Gli1-Cre^ERT^; Ddr2^fl/fl^* mice. Loss of *Ddr2* in *Gli1*-expressing cells faithfully recapitulated the cranial phenotype of *Ddr2^slie/slie^* mice including the presence of open frontal sutures (analogous to the metopic suture in humans) and thinning of frontal bones. This phenotype also mimics the open fontanels seen in SMED, SL-AC patients. These results are consistent with a function for *Ddr2* in Gli1+ sutural cells to regulate osteoblast differentiation during intramembranous bone formation.

Intriguingly, our lineage tracing studies identified *Gli1-Cre^ERT^*-induced tdTomato labeling beyond previously reported sites in cranial sutures, long bone growth plates and metaphyses.^6,32,41^ Specifically, this *Cre* also marked chondrocyte lineages and co-localized with *Ddr2* in cranial base synchondroses. Consistent with this distribution, tamoxifen treated *Gli1-Cre^ERT^; Ddr2^fl/fl^* mice had cranial base defects that were similar in type and magnitude to those seen in *Ddr2^slie/slie^* mice. Since it has been proposed that changes in the cranial base can indirectly affect suture formation, we conducted a second conditional knockout of *Ddr2* in chondrocytes using *Col2a1-Cre* that would be expected be preferentially active during endochondral bone formation in synchondroses, but not during intramembranous bone formation in sutures (Fig.3). The synchondroses of *Col2a1-Cre; Ddr2^f/f^* and *Gli1-Cre^ERT^; Ddr2^fl/fl^* mice were similar. However, *Col2a1-Cre; Ddr2^f/f^* mice had normal frontal sutures. Therefore, the contribution of *Ddr2* to endochondral bone formation in the synchondrosis is independent from its function in cranial sutures.

An unexpected finding was the observed thinning of frontal bones in *Col2a1-Cre; Ddr2^f/f^* mice. This thinning, which was first detected in *Ddr2^slie/slie^* and *Gli1-Cre^ERT^; Ddr2^fl/fl^* mice, would not normally be expected at a site of intramembranous bone formation that does not involve substantial synthesis of type 2 collagen. However, interpretation of this result is confounded by earlier studies showing endogenous *Col2a1* expression and *Col2a1-Cre* activity can be detected in calvarial osteoblasts and suture mesenchymal cells^42,43^. It is therefore possible that this *Cre* could inactivate *Ddr2* in a select population of *Col2a1*-expressing cells in intramembranous bone, thereby explaining the reduced frontal bone thickness in *Col2a1-Cre; Ddr2*^*fl*/fl^ mice. However, the absence of a posterior frontal suture defect *Col2a1-Cre; Ddr2*^*fl*/fl^ mice argues against Ddr2 having a function in *Col2a1*-expressing cells necessary for suture formation.

Fibrillar type II collagen, the predominant collagen species of cartilage matrix and one of ligands for DDR2, is important for normal chondrocyte proliferation as well as maintenance of the structural integrity and cellular organization of developing tissues.^44–48^ In this study, we evaluated type II collagen distribution in synchondroses using immunofluorescence and showed uniform staining throughout the cartilage matrix of wild type mice. However, Ddr2-deficient mice showed a non-homogenous staining pattern, predominantly in pericellular space adjacent to chondrocytes. Interestingly, similar results were observed in *Dmm/Dmm* mice harboring a mutation in the C-propeptide globular domain of type II collagen (*Col2a1*)^45^ as well as in mice containing an R992C mutation associated with spondyloepiphyseal dysplasia.^37^ These mutations cause defects in the assembly and folding of type II collagen and alter collagen distribution in cartilage ECM.^37,44,45^ Mutant mice are dwarf, have a shortened vertebral column and small rib cage as well as craniofacial defects characterized by skull dysmorphogenesis, reduced skull length, reduced cranial base length, cleft palate, and short mandible. These phenotypes are similar to those seen in *Ddr2*-deficient mice. While determining DDR2 regulation of various aspects of collagen fibrillogenesis is beyond the goal of this study, our findings are in agreement with previous studies showing abnormal collagen organization and/or crosslinking in *Ddr2*-deficient long bone growth plates, heart, postnatal testicular tissues and tumor-associated stroma.^32,49–51^ In addition, our previous characterization of *Ddr2* function in tooth development showed atypical collagen fibers in periodontal ligaments surrounding tooth roots in *Ddr2*^slie/slie^ mice.^31^ Most importantly, the altered distribution of cartilage matrix was also reported in a SMED, SL-AC patient.^52^ Histological analysis of resting zone of costochondral cartilage from a SMED, SL-AC patient showed abnormally distributed chondrocytes with noticeable dark staining in surrounding matrix filling lacunar space.^52^ The altered distribution of cartilage matrix highlights the similarity between histopathological features of the patient samples and our results in *Ddr2*-deficient mice, and provides important insights into potential mechanisms underlying *Ddr2* regulation of cartilage extracellular matrix.

The collagen-rich ECM maintains the structural integrity and cellular organization of developing tissues, controls the spatial and temporal response to growth factors and can activate several signal transduction pathways.^19,53,54^ We hypothesized that the defect in type II collagen distribution seen *Ddr2*-deficient mice could impact the ability of cells to maintain their spatial organization and polarization and disrupt cellular activities that underlie proliferation and differentiation of hypertrophic chondrocytes in the cranial base. Along these lines, we found that *Ddr2*-deficient synchondroses had an ectopic hypertrophic zone, disrupted chondrocyte polarization, and altered organization of Golgi apparatus, as shown by the asymmetric distribution of the Golgi marker, GM130. GM130 is a peripheral membrane protein tightly bound to the cytoplasmic face of the Golgi apparatus, the central organelle for secretory protein trafficking and cargo modification. Interestingly, a similar disruption of cell polarity was observed in mice mentioned above, which contain a R992C mutation in type II collagen.^37^ GM130 is required for maintaining Golgi structure, spindle assembly, cell cycle progression, and cytoskeletal dynamics.^55–59^ and disruption of GM130 in mitosis impairs spindle assembly and cell division.^55^ This may explain the underlying deficient chondrocyte proliferation seen in *Ddr2*-deficient synchondroses, although these mechanisms await elucidation.

Significant similarities and differences are apparent when results of the present study are compared with our previous work on Ddr2 functions in the appendicular skeleton^32^. At both sites, Ddr2 was localized to resting and proliferative zone cartilage of synchondroses or long bone growth plates, respectively, where co-localization with the Hh intermediate, Gli1, was observed. Lineage tracing at both sites suggested that Ddr2 positive cells could serve as progenitors for hypertrophic chondrocytes, osteoblasts, and osteocytes of endochondral bone. Consistent with this cartilage localization, inactivation of *Ddr2* in Gli1-positive progenitors (*Gli1CreERT*) or in chondrocytes (Col2a1Cre) inhibited endochondral bone formation in synchondroses and tibial growth plates and this inhibition was related to inhibition of chondrocyte proliferation rather than an increase in apoptosis. Type II collagen matrix organization and chondrocyte alignment was also disrupted at both sites although this phenotype was more severe in the synchondrosis where chondrocytes formed an ectopic hypertrophic zone. Ddr2 functions in endochondral bone formation likely explain the dwarf phenotype, reduced A-P skull length and shortened snout of Ddr2 deficient mice and as well as the dwarfism and the depressed facial features of SMED, SL-AC patients. In mice, Ddr2 was also expressed adjacent to intramembranous bone in the periosteum of calvaria and tibial cortical bone as well as cranial sutures. However, important differences in intramembranous bone formation were seen when comparing these two sites. In the cranial vault, Ddr2 deficiency inhibited frontal suture formation and reduced frontal bone thickness. In contrast, Ddr2 deficiency had little effect on thickness of tibial cortical bone, which also forms through an intramembranous process^32^. The basis for these differences is not clear. On one hand, differences in rates of formation of calvarial versus cortical bone may mask Ddr2-related activity in cortical bone. Alternatively, they may reflect inherent differences between these two tissue sites. Functions of Ddr2 in cortical bone may be more apparent in regeneration models which involve more rapid bone formation than is seen during development. As we recently showed, Ddr2 deficiency severely disrupts intramembraneous bone healing of subcritical-sized calvarial defects^60^. It is not known if healing of related cortical bone defects would also be disrupted.

In summary, this study establishes a critical function of *Ddr2* in skeletal progenitor cells, including GLI1+ cells, and chondrocytes to control bone formation and cartilage growth during postnatal growth of the craniofacial skeleton and advance our understanding of the roles of cell-matrix interactions in craniofacial morphogenesis. Our findings provide further insight into the cellular basis of cranial defects in SMED SL-AC patients as well as evolutionary changes in craniofacial structure related to altered DDR2 activity.

## Materials and Methods

### Mice

*Smallie* mice (*Ddr2*^slie/slie^) described in ref.^29^ were obtained from Jackson laboratory (stock no. 008172). *Ddr2^fl/fl^* mice with *loxP* sites flanking coding exon 8 of *Ddr2* gene, *Ddr2-LacZ* mice and *Ddr2^mer-iCre-mer^* mice harboring a MerCreMer cassette knocked in-frame into exon 2 of the *Ddr2* locus were previously described.^31,32^. For lineage analysis, *Rosa26-CAG-loxP-stop-loxP-tdTomato* mice (Ai14: R26R-tdTomato)^61^ were crossed with *Ddr2^mer-iCre-mer^* heterozygous mice to generate *Ddr2^mer-iCre-mer^*; Ai14: R26R-tdTomato mice. Tamoxifen was administered as previously described.^32^

To investigate the role of *Ddr2* in craniofacial development, we generated conditional knockout mice by crossing *Ddr2^fl/fl^* mice with tissue specific Cre driver mouse lines: *Col2a1-Cre* (gift from Dr. Ernestina Schipani, U Michigan);^35^ *Gli1-Cre^ERT6234^* (gift from Dr. Yuji Mishina, U. Michigan);^62^ *OC-Cre* (gift from Dr. Kurt Hankenson, U. Michigan).^36^ Mice used in our experiments were analyzed on C57BL/J6 background, except *Col2a1-Cre; Ddr2^fl/fl^* mice were analyzed on mixed background. All mice were housed under a 12-hour light cycle in compliance with the Guidelines for the Care and Use of Animals for Scientific Research. All protocols for mouse experiments were approved by the Institutional Animal Care and Use Committee of the University of Michigan. Genotyping of *Ddr2^mer-iCre-me^, Col2a1-Cre, Gli1-Cre^ERT^, OC-Cre* and Ai14: R26R-tdTomato mice was performed using PCR primers, as previously described.^35,36,61,62^ The genotyping of *Ddr2^slie/slie^* mice was performed using qRT-PCR with TaqMan probes on an ABI 7500 (Applied BioSystems).^28^

### Morphometric analysis of skulls

The craniofacial skeleton was examined by whole-mount skeletal staining as described previously.^63^ For microcomputed tomography analysis, 10 skulls from 3-month-old male and female mice were harvested and fixed in 10% formalin overnight at 4°C. Using microCT Scanco Model 100 (Scanco Medical), mouse skulls were scanned and reconstructed with voxel size of 12 μm, 70 kVp, 114 μA, 0.5-mm aluminum filter, and integration time of 500 ms. For craniofacial characterization, skull scans in VFF files were reoriented and cropped in MicroView software version 2.5.0 (see orientation in **Supplemental Fig.1**) and linear craniofacial measurements were made on 2D reoriented CT images using ImageJ software (version 1.51). We conducted skull linear measurements using previously published craniofacial landmarks.^33^ Thickness of cranial bones was measured on 2D reoriented CT images using a defined position on orthogonal views of MicroView software. Mid-bone regions (highlighted in blue and orange colors in **Fig. 1b**) were selected to measure the thickness the frontal and parietal bones on the coronal (frontal) plane, where right and left measurements were taken around frontal and sagittal sutures for frontal and parietal bone, respectively, and the average of two measurements were reported. Thickness measurements of the occipital bone were made on the sagittal plane as shown in **Fig. 1b,f**. Measurements for cranial base synchondroses were made on the mid sagittal plane, where synchondrosis width was determined by drawing a straight line in between chondro-osseous junctions with flanking bones and synchondrosis height determined by measuring the distance between two parallel lines tangential to the ventral and dorsal surfaces of synchondroses (**Fig. 2a**). The length of cranial base bones was quantified by measuring the distance between two lines defining anterior and posterior borders of individual cranial base bones. All linear measurements were made on reoriented 2D μCT images using straight-line function in ImageJ.

### Histology and Immunostaining

The whole skulls were fixed in 4% paraformaldehyde (PFA) for 48 h at 4°C. Specimens were decalcified in 10% ethylenediaminetetracetic acid (EDTA) (pH 7.2) and were then processed for paraffin embedding. Specimens were sectioned at 5 μm, deparaffinized and hydrated in ethanol series (100%, 95%, 70%) and in distilled water. For histological analysis, the sections were stained with hematoxylin and eosin according to the standard procedures. For immunofluorescence, sections were subjected to heat-induced antigen retrieval using 1X Diva Decloaker (Biocare medical) following the manufacturer’ instructions, washed with 1X PBS and then incubated with blocking buffer containing 5-10% normal donkey serum, 1% bovine serum albumin (BSA), 0.01% tween in 1X BSP for 1h at room temperature in a humid box. After blocking, the sections were incubated at 4°C overnight, with the following primary antibodies: anti-DDR2 (LS B15752, 1:200), Anti-COL2 (1:100, ab34712, Abcam); anti-COL10 (1:100, ab58632, Abcam); Anti-COL1 (1:100, AB765P, Millipore Sigma); Anti-GM130 (1:100, 610822, BD Biosciences); anti-IBSP (1:100). Anti-IBSP antibody was generated from a GST fusion protein containing amino acids 8-324 generated in the project laboratory. This was used as antigen for antibody production in rabbits (Harlan Laboratories). For Anti-GLI1 (1:100, NBP1-78259, Novus biological), sections were retrieved using citrate buffer (target retrieval solution, Dako) heated to 60 °C for 1 h. The slides were rinsed three times with 1X PBS, and the coverslip was mounted using ProLong™ Gold Antifade Mountant with DAPI (Life technologies) for cell nuclei staining. The sections were then imaged with a Nikon Eclipse 50i microscope and an Olympus DP72 camera.

### LacZ (β-galactosidase) Staining

LacZ staining of heterozygous Ddr2-LacZ (Ddr2^+/LacZ^) skulls was performed according to standard protocols.^64^ Samples were fixed in 2% paraformaldehyde and 0.2% glutaraldehyde in 0.1 M phosphate buffer pH 7.3 plus 5 mM EGTA and 2 mM MgCl_2_ at 4 °C. For the whole-mount staining, samples were rinsed after fixation 3 times in 1X PBS plus MgCl_2_, and then incubated at 37 °C overnight in a freshly prepared X-gal solution containing an X-gal substrate (UltraPure X-Gal, Invitrogen), 2mM MgCl_2_, 5mM potassium ferricyanide (III) (702587, Sigma), 5mM potassium hexacyanoferrate (II) (P3289, Sigma), 0.01% sodium deoxycholate, 0.02% NP-40. The samples were then visualized using a dissection microscope (Nikon SMZ 745T), and images were captured using a Nikon DS-fi1 camera. The wild-type littermates were used as controls. For frozen sections, samples were decalcified with 20% EDTA (pH 7.2) for up to 2 weeks according to the mouse age, rinsed in 1X PBS, then placed in 30% sucrose in 0.1M phosphate buffer with 2mM MgCl_2_ at 4 °C for overnight. Samples were then embedded in optimal cutting temperature compound (Tissue-Tek) and cryosectioned at 12 μm thickness at −20 °C, mounted on glass histological slides (Fisherbrand™ ColorFrost™ Plus). For LacZ staining of frozen sections, sections were post-fixed in 0.2% PFA in 0.1 phosphate buffer pH 7.2 for 10 min on ice, washed 3 times in MgCl_2_-containing PBS, and stained with an X-gal substrate (UltraPure X-Gal, Invitrogen), 2mM MgCl_2_, 5mM potassium ferricyanide (III) (702587, Sigma) and 5mM potassium hexacyanoferrate (II) (P3289, Sigma) overnight at 37°C. The sections were washed 3 times in 1X PBS followed by distilled water, counterstained with Vector Nuclear Fast Red staining, and dehydrated through ethanol series and xylene and mounted with an Acrytol mounting medium (Leica).

### Proliferation and Apoptosis Assay

To assay cell proliferation, mice were injected intraperitoneally with 5-ethynyl-2’-deoxyuridine and sacrificed 4h after injection. EdU-labeled cells were detected using Click-iT® EdU Alexa Fluor® 488 Imaging Kit (Invitrogen, # C10337). Briefly, after deparaffinization and hydration, the tissue sections were incubated with Click-it reaction mixture for 30 min in a dark humidified chamber. The sections were washed 3 times with 1X PBS for 2 minutes each, mounted with ProLong™ Gold Antifade Mountant with DAPI (Life technologies) and imaged with a Nikon Eclipse 50i microscope and an Olympus DP72 camera. To assay cell apoptosis, we performed terminal deoxynucleotidyl transferase dUTP nick end labeling (TUNEL) assay according to the manufacturer’s instructions (FragEL™ DNA Fragmentation Detection Kit, Colorimetric-Klenow Enzyme, Calbiochem). Briefly, after deparaffinization and rehydration, the tissue sections were rinsed in 1X tris-buffered saline (TBS), permeabilized with Proteinase K and the endogenous peroxidase activity was blocked with 3% hydrogen peroxide for 5 minutes at room temperature. The sections were then incubated with Klenow Labeling reaction mixture in a humidified chamber at 37 °C for 1.5 h. For labeling detection, the sections were incubated with peroxidase streptavidin conjugate and subsequently with DAB solution. The sections were counterstained with methyl green solution and imaged with a Nikon Eclipse 50i microscope and an Olympus DP72 camera.

### Statistical Analysis

The graphs and statistical analysis were performed in the GraphPad Prism software (version 6.0e, La Jolla California USA). Mouse studies used an N = 10 based on power analysis of data from our previous study with Ddr2-deficient mice^28^ where we estimate a minimum of 8 animals/group will be required to detect a power of .80 (95% CI, estimated effect size of η2 >0.40). All values were presented as mean ± S.D. Unpaired, two-tailed Student’s *t* test was used to analyze the difference between the two experimental groups. * *P* < 0.05; ** *P* < 0.01, *** *P* < 0.001, **** *P* < 0.0001; n.s. not significant.

## Acknowledgements

This work was supported by a scholarship from the Ministry of Higher Education and Scientific Research, Libyan Transitional Government (FFM), a scholarship from King Saud University (AB), NIH/NIDCR grants DE11723, DE029012, DE029465, Department of Defense grant PR190899, research funds from the Department of Periodontics and Oral Medicine, University of Michigan School of Dentistry (to RTF) and the Michigan Musculoskeletal Health Core Center ((NIH/NIAMS P30 AR069620).

## Author Contributions

FM, CG and AB conducted experiments; RC and BG developed *Ddr2^mer-iCre-mer^* mice; NO assisted with lineage tracing studies; FM, CG and RTF conducted all data analysis; FM and RTF wrote the manuscript. All authors critically reviewed the manuscript.

Supplementary information accompanies the manuscript.

---

**Supplementary Fig. 1:**
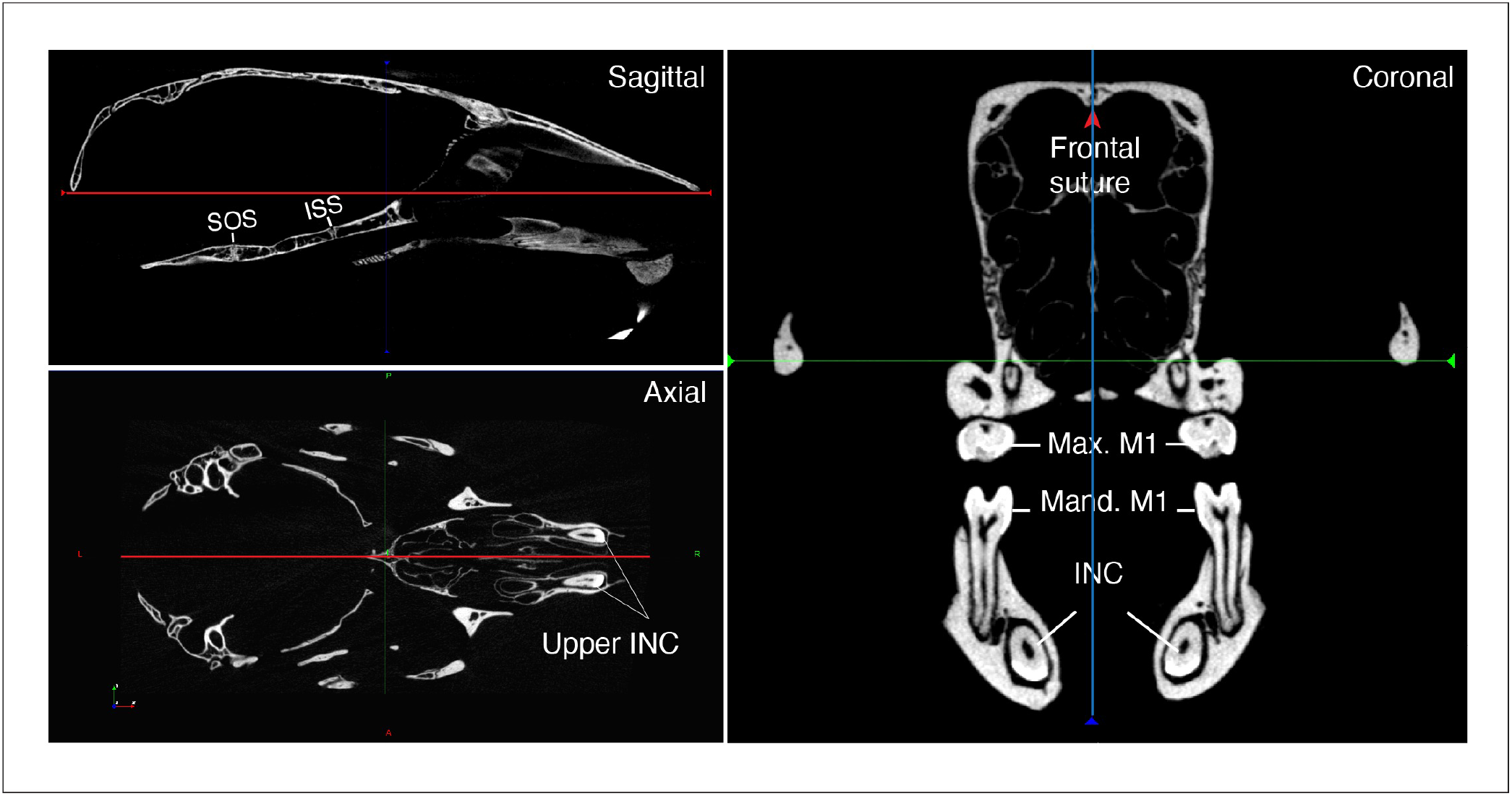
Orientation of skulls in sagittal, axial and coronal planes. Max: Maxillary; Mand: Mandibular; M1: first molar; INC: Incisor; ISS: Intersphenoid synchondrosis; SOS: Spheno-occipital synchondrosis.

**Supplementary Fig. 2:**
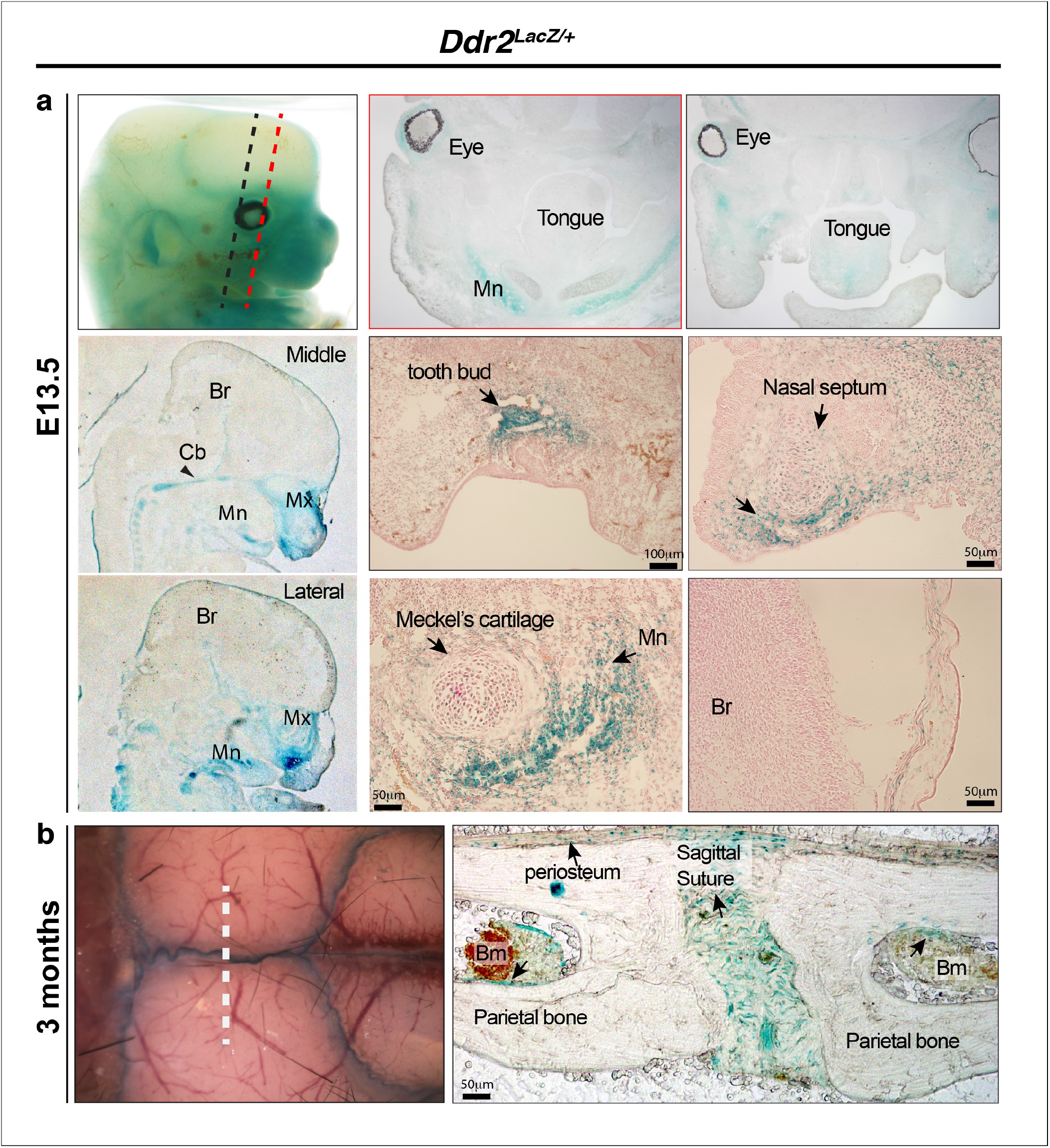
Ddr2-LacZ localization in craniofacial skeleton. **a**, In *Ddr2^+/LacZ^* embryos harvested at E13.5, intense X-gal staining can be seen in the maxilla (Mx), mandible (Mn), cartilage primordia of cranial base (Cb, black arrowhead), nasal septum and tooth buds. No staining was detected in the brain (Br) (top row, whole mount X-gal staining; middle and bottom row, X-gal staining on frozen embryos). **b**, Whole mount and frozen section of calvariae from 3-month old mice show intense X-gal staining in cranial sutures, and also in periosteum, and the lining of bone marrow inside cranial bones.

**Supplementary Fig. 3:**
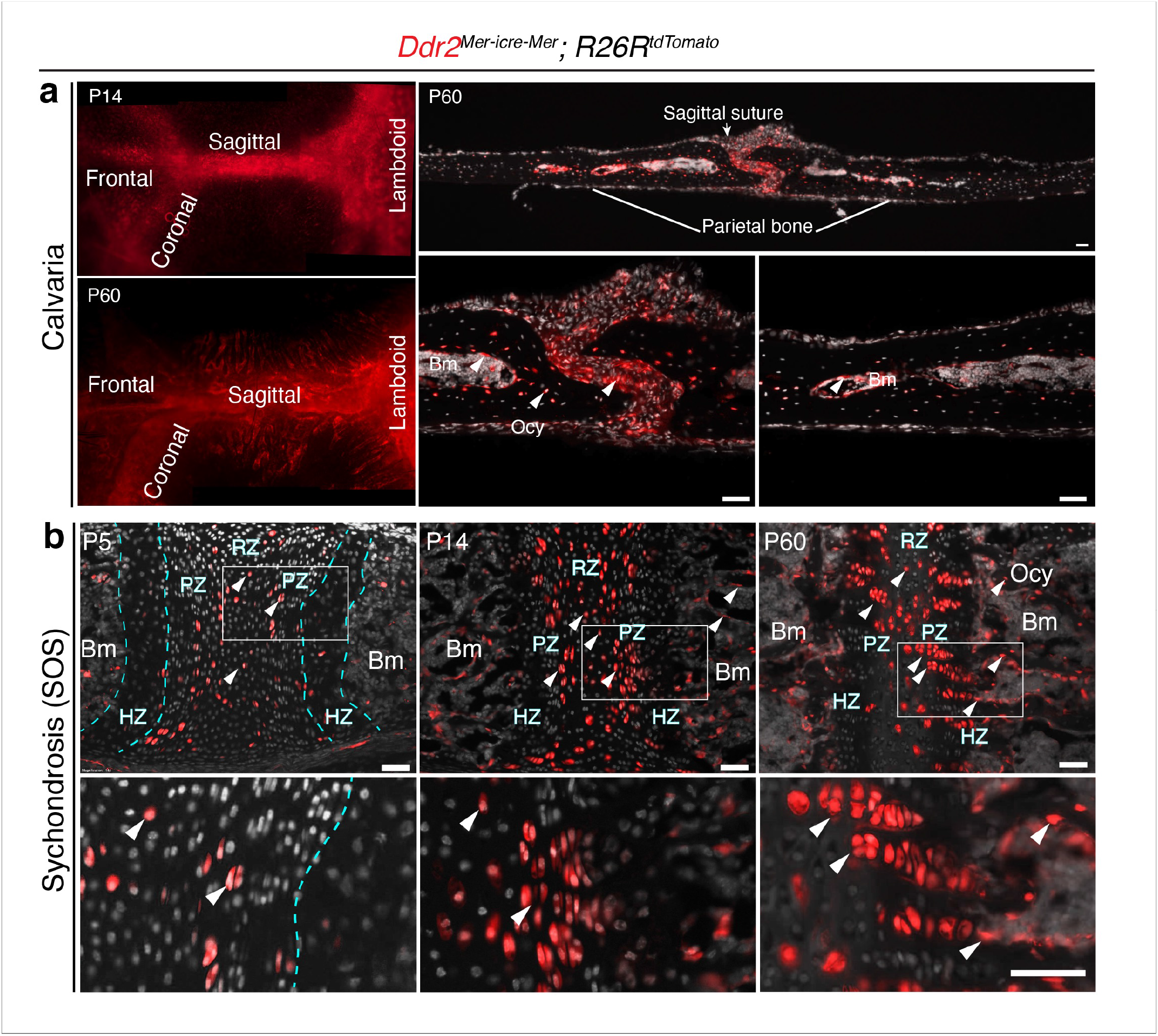
*Ddr2^mer-iCre-mer^* induced recombination in cranial sutures and cranial base synchondrosis. Neonatal *Ddr2^Mer-icre-Mer^*;Ai14^CAG-tdTomato^ mice were treated with tamoxifen as described in Fig. 5. **a**, Whole mounts (left) at P14 and P60 show *Ddr2 tdTomato* labeling in all cranial sutures: frontal, sagittal, coronal and lambdoid sutures (left). Cryosections (right) show distribution of *tdTomato-labelled* cells at P60 in the suture mesenchyme, bone marrow lining cells and osteocytes. Scale bar: 50μm. **b**, Cryostat sections of the cranial base spheno-occipital synchondrosis (SOS) shows *tdTomato* labeling initially in resting and proliferative chondrocyte zones (P5). At later times (P14, P60), progeny of *Ddr2*-positive cells form single or two column clones along the axis of cranial base growth extending into the hypertrophic zone and osteocytes. Scale bar: 50μm.

**Supplementary Fig. 4:**
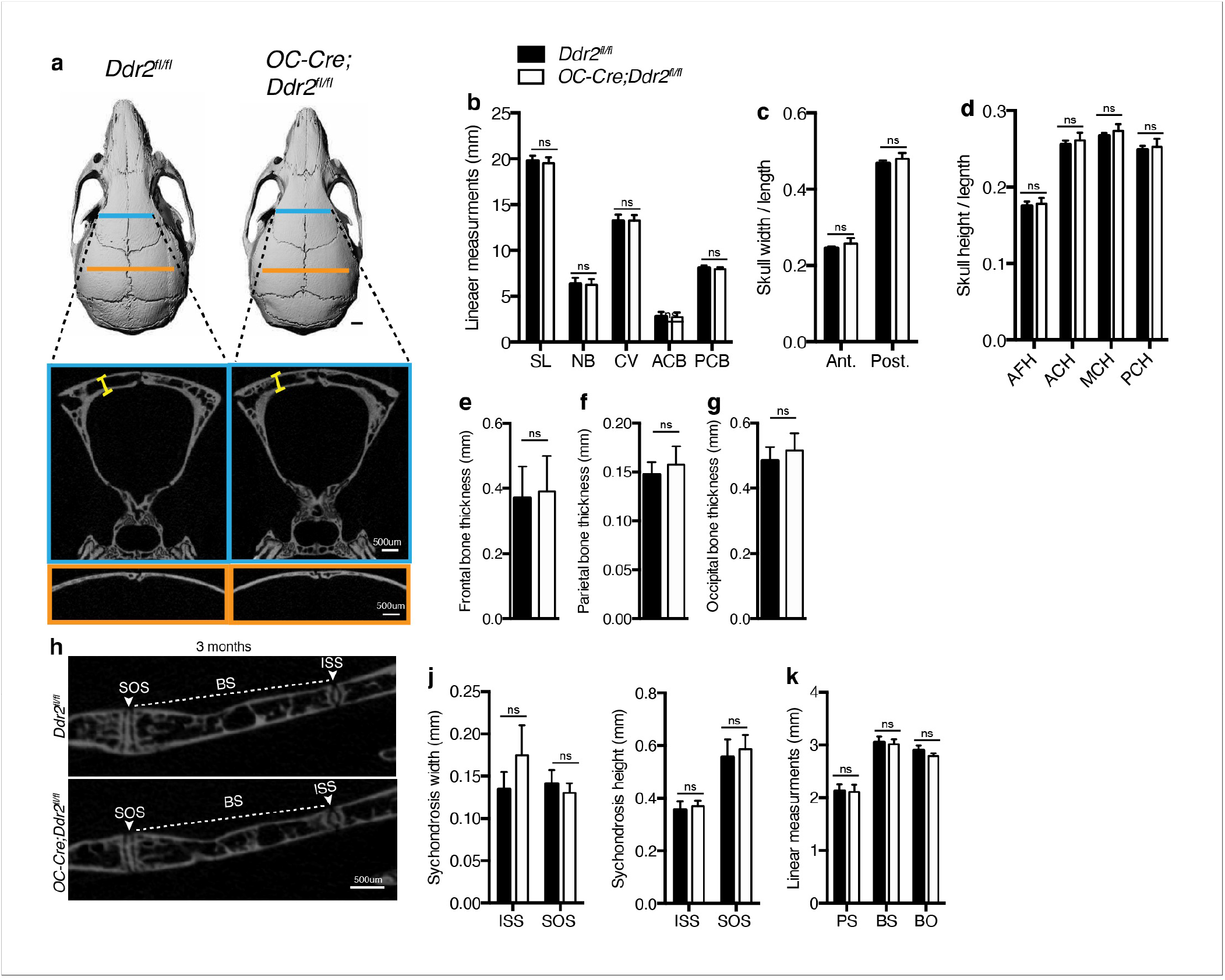
*Ddr2* loss in mature osteoblasts using OC-Cre did not result in craniofacial abnormalities. **a-e**, Quantification using μCT scans shows no difference between *Ddr2^fl/fl^* and *Ocn-Cre; Ddr2^fl/fl^* skull length, width and height at age of 3 months. Scale bar: 1mm in a. **e-g**, quantification showing no difference in frontal, parietal and occipital bone thickness. **h-k**, Quantification using μCT scans showing no changes in cranial base synchondroses or associated bones. Data are presented as mean ± SD. (*n*=10). ns, not significant, two tailed- unpaired *t* test.

**Supplementary Fig. 5:**
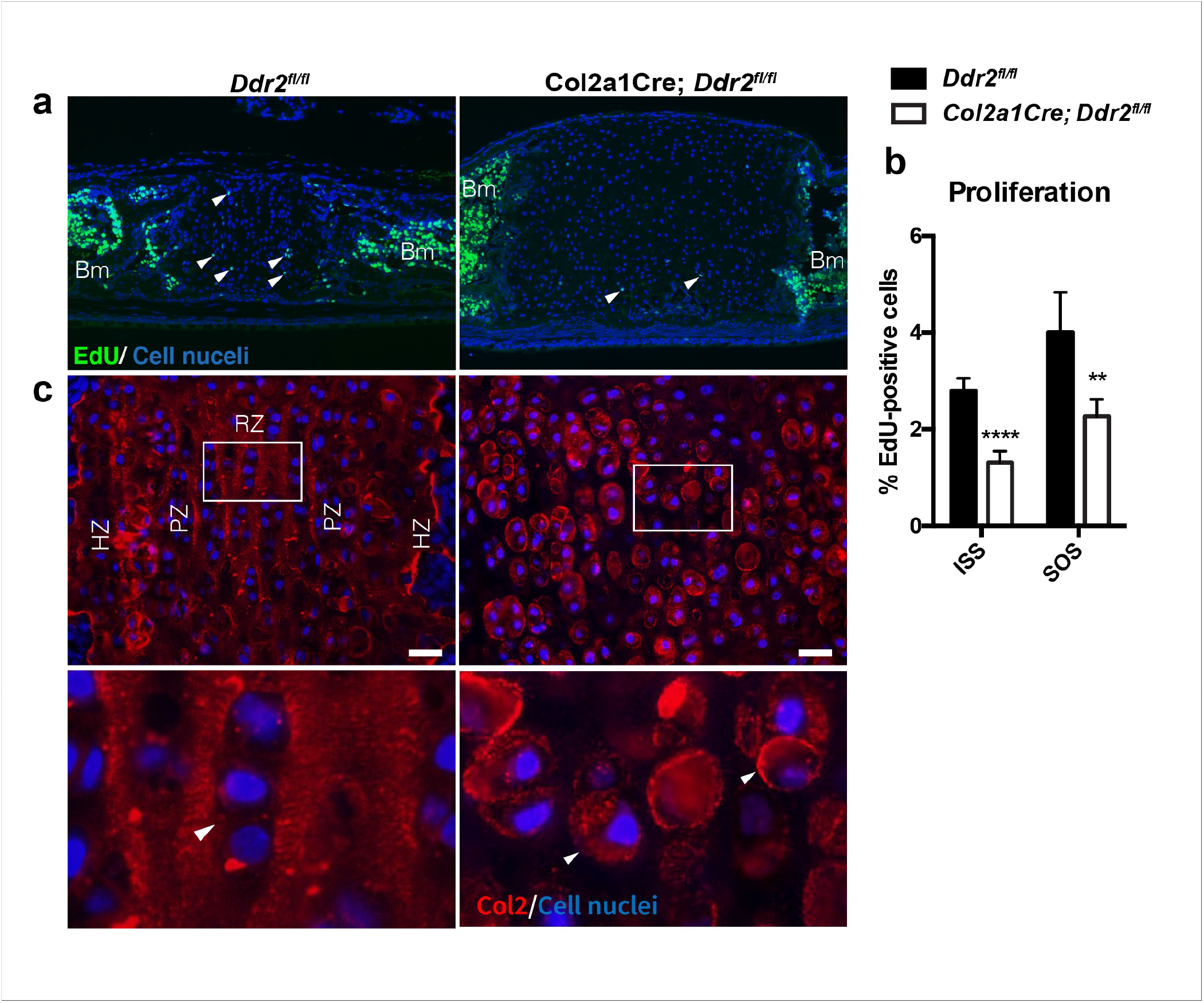
Synchondroses in *Col2a1Cre:Ddr2ff* mutants exhibited deficient chondrocyte proliferation and abnormal type II collagen distribution. **a**, EdU staining (green) showed a significant reduction of chondrocyte proliferation in *Col2a1-Cre; Ddr2^fl/fl^* synchondrosis. **b**, Bar graph shows quantification of EdU-positive cells in ISS and SOS. **c**, COL2 immunostaining shows altered type II collagen matrix in *Col2a1-Cre; Ddr2^fl/fl^* mice. n= 3 mice, \*\**P*<0.01, \*\*\*\**P*<0.0001. Scale bar: 50 μm in (a), 20 μm in (c).

